# Environmental filtering of life-history trait diversity in urban populations of *Arabidopsis thaliana*

**DOI:** 10.1101/2022.10.03.510679

**Authors:** Gregor Schmitz, Anja Linstädter, Anke S. K. Frank, Hannes Dittberner, Jessica Thome, Andrea Schrader, Karl-Heinz Linne von Berg, Andrea Fulgione, George Coupland, Juliette de Meaux

## Abstract

1. The challenges to which plants are exposed in urban environments represent, in miniature, the challenges plants face as a result of global environmental change. Hence, urban habitats provide a unique opportunity to assess whether processes of local adaptation are taking place despite the short temporal and geographical scales that characterize the Anthropocene.
2. We quantified the ecological diversity of urban habitats hosting *A. thaliana* populations. Using plant community indicators, we show that these patches differ in their levels of soil nutrient content and disturbance. Accordingly, plants in each patch displayed a range of flowering time, size and fitness.
3. Using a deep sampling approach coupled with reduced genome-sequencing, we demonstrate that most individuals can be assigned to a limited set of clonal lineages; the genetic diversity of these lineages is a sample of the diversity observed in western European populations of the species, indicating that established urban populations originate from a broad regional pool of lineages.
4. We assessed the genetic and phenotypic diversity of these lineages in a set of common garden experiments. We report marked genetic differences in life-history traits, including time of primary and secondary dormancy as well as of flowering. These genetic differences in life-history traits are not randomly distributed but sorted out by ecological differences among sites of origin.
5. **Synthesis:** Our study shows that the genetically diverse phenology of a regional *A. thaliana* gene pool is not randomly distributed but filtered by heterogeneity in the urban environment. To out knowledge, this report is the first to show a pattern indicative of environmental filtering enhancing local genetic adaptation within urban environments. We conclude that environmental filtering helps maintain functional diversity within species.

## 1. Introduction

Urban environments provide untapped opportunities to link ecological and phenotypic diversity over the short temporal and spatial scales that define the Anthropocene (Johnson & Munshi-South, 2017; Szulkin et al., 2020). Understanding how plant functional variation is optimized within local urban environments can help understand the potential of species to cope with upcoming global changes.

Compared to that in neighboring rural areas, the climate in urban environments is generally characterized by increased temperature and reduced moisture, and thus a higher drought likelihood (Lambrecht et al. 2016). In addition, small-scale differences within urban microclimates are observable, depending on urban geometry and construction materials (Chatzidimitriou & Yannas, 2015). Urban microhabitats differ in edaphic characteristics: soils vary considerably in their developmental stage, depth, compaction, and surface transformation (Foti, 2017; Johnson & Munshi-South, 2017; Tresch et al., 2018). Shallow, sealed soils with a low capacity for water storage and low nutrient content typify the so-called urban grey spaces associated with built urban elements, such as sidewalks. They occur in close vicinity to nutrient-rich urban green spaces, often with eutrophic or even contaminated soils, which results in huge variation in soil-mediated plant resources over small spatial scales (Bonthoux et al., 2019; Gilbert, 2012; Vega & Küffer, 2021). Cities also host numerous ruderal sites, which are exposed to frequent and often severe disturbances, fragmentation, and high turnover rates among populations (Dubois & Cheptou, 2017; Guo et al., 2018). In addition, human-driven disturbances, such as weeding or mowing, may shorten the time windows for growth and reproduction, and thus modulate further the disturbance patterns.

Major life-history decisions, such as the timing of germination and the timing of flowering, shape the timing of plant phenological transitions and are critical for resource acquisition and reproductive success. They have been shown to contribute to the persistence of populations in heterogeneous urban settings (Liu et al., 2021; Nord & Lynch, 2009). Plant phenology is plastic. It can be adjusted locally in response to the environment because the life-history traits underlying it are modulated both by nutrient resources and by environmental cues, such as temperature and photoperiod (Vidal et al., 2014). For example, flowering is accelerated by winter temperature and resource stress and by high temperatures in spring, and delayed by the shortening days of the fall (Andrés & Coupland, 2012; Li et al., 2019). Plant phenology thus shifts in response to environmental cues that are modified by climate change (Chmielewski & Rötzer, 2001; Piao et al., 2019; Primack & Miller-Rushing, 2012).

The genetic composition of the population can also be tuned by local selective pressures acting on the genetic basis of phenological variation. Adaptation to local environments has been observed experimentally in many organisms (Kawecki & Ebert 2004; Leimu & Fischer 2008). Clearly, adaptation via new genetic mutations is unlikely within urban settings because urban populations are small, fragmented, and prone to frequent extinction(Orr, 2005). However, if the pool of potential migrant genotypes is sufficiently diverse, then individual genotypes may establish or not depending on their genetic makeup, which results in local adaptation by environmental filtering. Indeed, patterns of local adaptation detected over small distances suggest that this kind of environmental filtering could happen in urban habitats (Halbritter et al., 2018; Lenssen et al., 2004; Robionek-Selosse et al., 2021).

In recent years, several studies have described phenotypic differences and genetic differentiation between individuals from urban and neighboring rural sites, consistently indicating that populations adapt rapidly to cities. These studies documented selection for novel optima in life-history traits, such as a late flowering time and decreased dispersal in holy hawksbeard (*Crepis sancta);* an early flowering time in common ragweed (*Ambrosia artemisiifolia*); and decreased cyanogenesis in white clover (*Trifolium repens*) (Cheptou et al., 2008; Gorton et al., 2018; Lambrecht et al., 2016; Santangelo et al., 2022; Thompson et al., 2016). Altogether, however, the amount of genetic variation in life-history traits, the distribution of this variation within the heterogeneous environments of cities, and its contribution to local adaptation in these environments remain unknown (Johnson & Munshi-South, 2017).

Urban environments in regions with temperate climates often host populations of short-lived herbaceous species, including the Brassicaceae species *Arabidopsis thaliana* (Bomblies et al., 2010; Herben et al., 2016; Huang et al., 2018). This species -- the initial model of plant molecular biology -- has become a powerful system for answering fundamental questions of ecology, evolution, and global environmental change (Exposito-Alonso et al., 2018; Fournier-Level et al., 2011; Takou et al., 2019; Vasseur et al., 2018). *A. thaliana*, which is native to large areas of Eurasia and Africa, has recently colonized many other parts of the world (Durvasula et al., 2017; Lee et al., 2017). The deep sampling of natural populations across various regions of its area of distribution has shown how environmental heterogeneity shapes the distribution of genetic diversity in *A. thaliana* (Castilla et al., 2020; DeLeo et al., 2020; Dubin et al., 2015, p. 20; Luo et al., 2015). Life-history trait adaptation appears to happen in concert across regional environments (reviewed in Takou et al. 2019). This includes adaptation in the timing of distinct phenological stages such as germination (Kronholm et al., 2012), or the onset of flowering (Ågren et al., 2017; Akiyama & Ågren, 2018; Le Corre, 2005), but also the tuning of other life-history traits such as the rate of rosette growth or leaf economic spectrum (Debieu et al., 2013; Sartori et al., 2019; Wieters et al., 2021).

In the present study, we determined the relative importance of phenotypic plasticity and genetic variation in shaping the phenology and fitness of eight spontaneously occurring urban populations of *A. thaliana*. We asked i) Are *A. thaliana* urban populations occupying ecologically diverse habitats? ii) Are *A. thaliana* populations genetically diverse? iii) Is this genetic diversity randomly distributed across habitats? By combining plant-community and trait-based ecological measurements with analyses of genetic diversity at genomic and phenological levels, we show that genotypes growing in urban habitats exposed to high resource limitations and increased disturbance levels tend to have traits tailored to local conditions.

## 2. Materials and Methods

### Study area

Our study focused on urban *A. thaliana* populations in the city of Cologne, situated in midwestern Germany (about 50.9°N, 7.0°E). Cologne is Germany’s fourth most populous city, with more than one million inhabitants (Ansmann et al., 2021). One of Germany’s warmest cities, it has a temperate oceanic climate with a mean annual temperature of 11.7 °C during the day and 6.3 °C at night, and a mean annual precipitation of 840 mm.

### Field surveys, phenology monitoring, and functional trait measurements

We surveyed eight *A. thaliana* populations within a 4.3 x 3.7 km area in southwestern Cologne. Study populations were selected based on their accessibility and on their isolation from other populations, with all habitat patches located at least 0.6 km from one another. Urban habitat patches where viable *A. thaliana* populations were observable included green spaces such as disturbed lawns; various habitats within gray spaces, such as sidewalks, wall tops, and bases; and vegetated roadsides (Figure 1a; Figure S1; Table S1). Depending on habitat characteristics, patch size ranged from 4 m^2^ for populations on wall tops to > 100 m² for populations in vegetated roadsides, and population size ranged from 15 to 1,000 individuals. All populations were studied throughout their growth period in 2017 (which extended between early March and the end of June), and revisited yearly during five subsequent growth periods (2018-2022) to assess population persistence. No field permissions were necessary for the collection of the plant samples in this study.

**Fig. 1:**
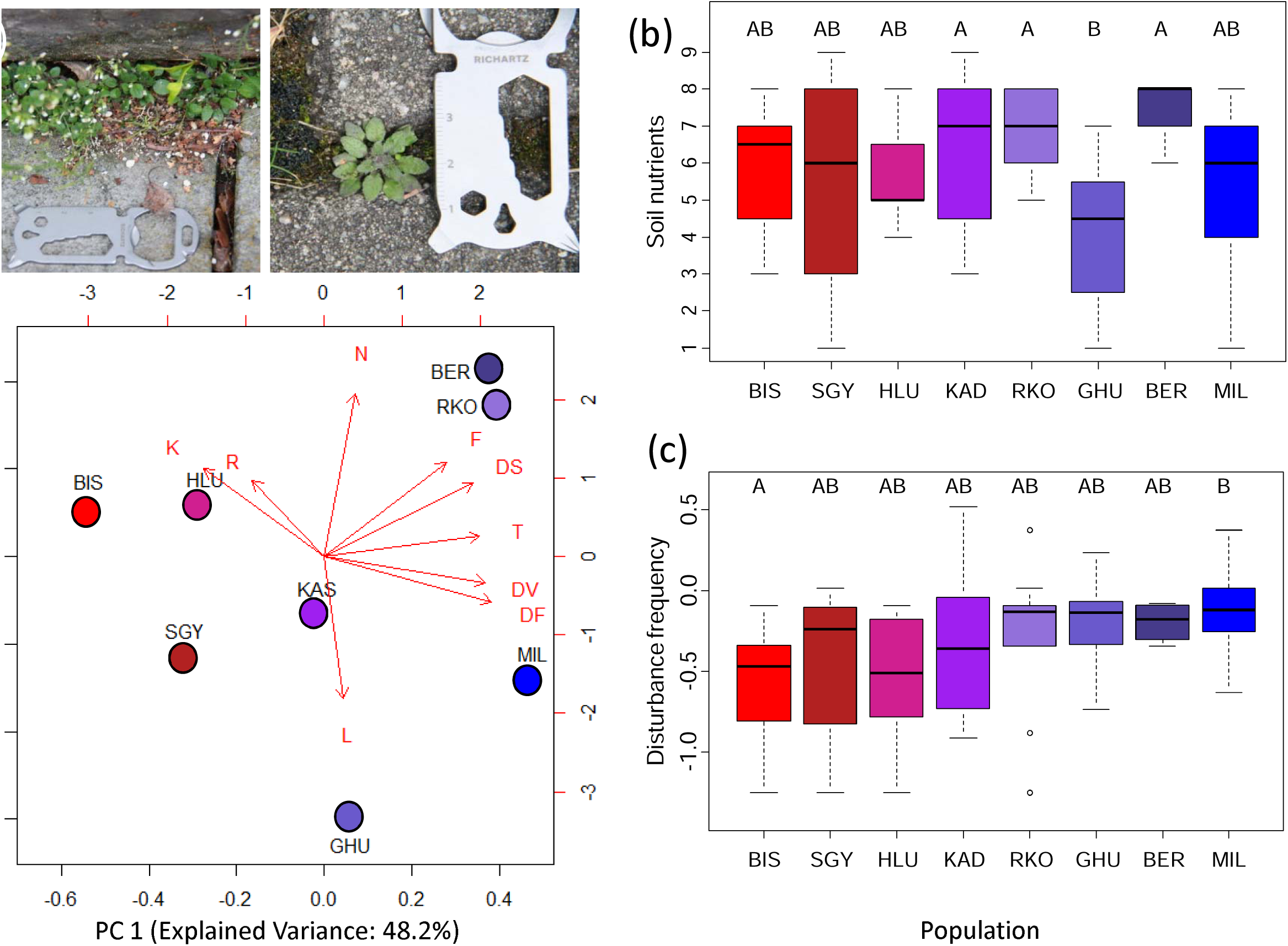
Characteristics of eight urban *Arabidopsis thaliana* populations and their habitat conditions in the city of Cologne, with (a) differences in size and phenological stage between *A. thaliana* from two adjacent habitats (both photographs from 2017-04-03); (b) differences between habitat patches based on plant species’ Ellenberg indicator values for soil nutrients and (c) the same based on plant species’ Herben indicator for disturbance frequency. Letters indicate significant differences at p<0.05. (d) Similarity of habitat patches based on a PCA with plant communities’ average indicator values for abiotic environmental conditions and disturbances. For indicator names, see Materials and Methods.

We recorded phenological stages and fitness-related plant functional traits of *A. thaliana* individuals distributed over the eight habitat patches. At each patch, we established 10 to 14 observation quadrats (10 x 10 cm), each at least 50-60 cm from one another. We marked from 1 to 3 *A. thaliana* individuals per quadrat (15-24 per habitat patch; 174 in total) and visited them twice a week during the study period to score their development. For all marked individuals, we recorded the day of bolting (i.e., when the inflorescence became visible) and, subsequently, the number of open flowers and the number of closed fruits. Rosette diameter and the number of rosette leaves were measured at the time of bolting. We used these observation data to derive five phenological stages and five fitness-related plant functional traits (“fitness proxies”; see Table S2). The selection of plant functional traits follows recommendations for the assessment of plant fitness (Gibson, 2014), while phenological stages were selected to represent life-history stages of the entire growth period, as recommended for herbaceous species (Huang et al., 2018; Nordt et al., 2021). Germination was generally not possible to determine *in situ*, except in populations growing on wall tops (SGY, KAD, HLU).

### Assessment of environmental conditions of habitat patches

Abiotic environmental conditions and characteristics of disturbance regime were recorded at the level of habitat patches, making use of the indicator function of plant communities (Kollmann & Fischer, 2003). We used species’ Ellenberg indicator values, EIVs (Ellenberg & Leuschner, 2010), as a proxy for abiotic conditions. EIVs, which express the optimal positions of Central European plant species along gradients of abiotic factors, provide information about species’ niche and habitat requirements (Diekmann 2003), in both rural and urban habitats (Fanelli et al. 2006). EIVs are expressed on an ordinal scale ranging from 1 to 9 and refer to light regime (L), temperature (T), and continentality of climate (K). Edaphic conditions are captured as soil moisture (F), pH (R), and nutrient availability (N). Likewise, species’ disturbance indicator values (DIVs) (Herben et al., 2016), were used to characterize the disturbance regimes of urban habitat patches. To this end, we extracted species-level indicators for disturbance severity (DS), disturbance frequency (DF), and disturbance effects on vegetation structure (DV) from the data compiled in Herben et al., 2016. DIVs are based on vegetation-plot records of the Czech flora, with a species’ DS defined as the mean disturbance severity of all vegetation classes weighted by its occurrence frequencies in these classes; DF calculated as the mean of logarithmic disturbance frequency of all vegetation classes weighted by a species’ occurrence frequencies in these classes; and DV calculated from vegetation structural parameters by summing covers and community-weighted means of all species’ growth height per vegetation class. We determined the composition of plant communities in autumn (late August and early September) 2017 by recording the presence of all vascular plant species in the herbaceous layer of the habitat patches. Based on species’ specific indicator values (see Table S4) and excluding *A. thaliana*, weighted average EIVs and DIVs were calculated for the plant community of each habitat patch.

### Assessment of genomic variation

At the end of the 2017 growth period, we harvested seeds from 5 to 17 individuals per population and used single seeds from each individual to produce progenies. For four of the eight habitat patches (BER, BIS, KAD, and MIL), we also included seeds from individuals sampled in the year prior to field data collection, referring to these as population samples (Table S3). In order to broaden our view of the genomic diversity of *A. thaliana* in Cologne, we also produced progenies from seeds collected at 30 additional habitat patches distributed throughout the sampling area but not included in the *in situ* monitoring (hereafter called scattered samples). Field collected seeds were grown in a randomized setting, under conditions simulating winter, i.e., in growth chambers with 10 h light at 18 °C / 14 h dark at 16 °C, 60 % humidity, for 6 weeks; followed by 8 h light/16 h dark at 4°C, 80 % humidity, for 6 weeks; and ultimately ripened in the greenhouse in spring-like conditions with 16 h light, at about 20-24 °C / 8 h dark at about 18-20 °C until harvest. Seeds for progeny were harvested using Aracons (Betatech, Gent, Belgium).

Three leaves per plant were sampled for DNA extraction, deep frozen, and homogenized using a Precellys Evolution homogenizer (Thermo Fisher Scientific, Waltham, MA, USA). DNA was isolated using the NucleoSpin® Plant II kit (Macherey-Nagel, Düren, Germany). Genomic DNA was prepared for RAD-Seq as described by (Dittberner et al., 2019) using 10 pools with 20 individually barcoded plant samples each, and sequencing was performed at the Max Planck Institute of Plant Breeding Research. The bioinformatics pipelines used for read mapping, and identification of each of the 12 genotypes present in the studied habitat patches, population structure and admixture are described in the supplementary information.

### Common garden experiments

For each identified genotype, two individuals were randomly selected and amplified in the greenhouse. Since they were raised in the same maternal environment, variance among genotypes can be assigned to genetic effects and variance among replicate lines of each genotype allows testing whether all maternal effects were effectively removed when growing parents in common garden conditions. We then took eight individuals from each of these two lines and grew them in growth chambers (Johnson Controls, Milwaukee, WI, USA) in three trials under different growth conditions. Growth conditions were chosen to identify genetic variation in the major pathways regulating flowering time: constant long days (16 h light at 20°C/8h dark at 18°C); constant short days (8 h light at 18°C day/16 dark at 16°C night); and short days/vernalization/long days (8 h day light at 18°C/16 h dark at 16°C for 5 weeks; 8h day light/16h dark at 4 °C for 4 weeks, 16 h day light at 20°C/8h dark at 18°C until ripening). The *A. thaliana* genotypes Col-0 was included as a reference in all experiments. Because we did not detect differences between replicated lines, we considered the number of replicates to be 2×8= 16 replicates/genotype. Replicates were sown in individual 6-cm-diameter pots that were watered regularly by flooding the trays. Flowering time was determined as the time span between sowing and the first flower to open petals.

Ripe seeds from the short-day/vernalization/long-day experiment were harvested and stored at room temperature for the germination assay. Germination rates were determined after three months to quantify primary dormancy (Baskin & Baskin, 2004). Seeds were sown on wet filter paper and incubated in closed microtiter plates. Before incubation under long-day conditions (16 h light at 20°C/8h dark at 18°C), seeds were incubated in the dark either at 4 °C for 7 days to quantify their capacity to germinate when dormancy is released, or at -21 °C for 4 d and at 35 °C for 4 d to quantify variation in secondary dormancy. Germination was scored after 10 days.

To quantify genetic variation under outdoor field conditions, we conducted a set of common garden experiments. For this, seeds from 4 independent replicate lines per genotype were sown directly at a density of 8 seeds per pot. In total, we planted 32 pots per genotype and randomized 9-cm-diameter pots at a field site in the botanical garden of Cologne University. Again, because we found no significant differences between replicate lines, we determined that the experiment was complete with 32 replicates per genotype. Seeds were sown in late summer (August 23) and in early autumn (September 20), mid-autumn (November 8), and late winter (February 26). Rather than being watered artificially, pots were put on a fabric that retained water after rainfall. The site was protected against birds and rabbits by a net. Plants were inspected for germination twice a week. Flowering time was scored when the first flower showed open petals. For fertility measurement, siliques were counted at the end of each plant’s life cycle.

Phenotypic data was analyzed using the glm function in R version 4.1.0 (R Core Team, 2022). We used one-way ANOVA with habitat patch as the fixed factor to analyze differences in environmental conditions (EIVs and DIVs of plant species co-occurring with *A. thaliana*); and in the phenological stages and fitness proxies of *A. thaliana* plants observed *in situ*. Measurements of flowering time in common garden experiments were analyzed with two-way ANOVA, using genotype and cohort or growth condition as fixed factors.

For germination assays under controlled indoor conditions, the number of germinants per Petri dish was analyzed with the R function zeroinfl (Package pscl; Zeileis et al., 2008) to correct for zero inflation. For germination rates in the outdoor common garden experiment, we proceeded as under controlled conditions, except that we used a quasipoisson distribution of error. Because the two replicate lines we used for each genotype never showed significant effects, the line effect was not included in any of the models. We used the function ANOVA with a Chi-squared test to determine the overall significance of effects. In all cases, the significance of pairwise differences was tested with a Tukey post-hoc test, using the function glht from the multcomp package in R (Hothorn et al., 2008). We used the Pearson coefficient of the cor.test function in R to quantify the correlation between phenotypes.

## Results

### Urban habitat patches of *A. thaliana* are ecologically diverse

The eight urban habitat patches differed significantly in environmental conditions related to climate, soil, and severity of disturbance. Indicator values for light (L), and edaphic conditions -- humidity (F), and soil nutrients (N) -- differed between sites (Figure 1b, supplemental Figure 1a, b; humidity (F): F_7,113_=2.855, p=0.00883; soil nutrients (N): F_7,93_=2.777, p=0.0114, light (L): F_7,130_=2.329, p=0.0285). Indicator values of the accompanying flora for herb-layer disturbance frequency (DF) and herb-layer structure-based disturbance (DV) differed between sites (Figure 1c, supplemental Figure 1c; DF: F_7,84_=2.897, p=0.0092; DV: F_7,84_=3.108, p=0.00577), whereas differences in herb-layer disturbance severity (DS) were not significant. The three values for disturbance were strongly and positively correlated with each other (range of r: 0.802 – 0.958; p<0.00172) and with the temperature indicator (range of r: 0.736 – 0.803; p<0.00632; Figure 1d; Figure S3, Table S4).

### Plants differ across habitat patches in their phenology and fitness

Across the eight urban *A. thaliana* populations, all phenological stages and plant functional traits measured *in situ* showed significant differences (p<2.43e-06; F=6.094, Table S6; Figure S4). These are exemplified in data on flowering time and rosette size (Figure S1a,b). The diverse traits describing phenological time points, the temporal spacing between phenological states, and the size of the plants also covaried positively (Figure S5).

### Community-level indicators of environmental variables explain part of the phenotypic differences between field sites

*In situ* observations of phenotypic differences between sampling sites were partly correlated with the differences in environmental characteristics: as expected, plants from sites with high mean values for N- and F-indicators, which reflect differences in nutrient and soil moisture levels, respectively, were on average larger than those from sites with lower N- and F-indicator values (p-values smaller than 0.05 for F: inflorescence length, maximal number of open flowers, rosette diameter; for N: rosette diameter at bolting, number of rosette leaves, and number of fruits; Figure S1c). Plants from sites with higher disturbance values bolted later than plants from other sites, but formed larger inflorescences bearing more fruits.

### Urban populations in Cologne consist of clones of diverse western European genotypes

In order to determine the extent of genomic variation and the history of population establishment, we genotyped 5-17 individuals per population as well as plants sampled from additional sites scattered across the study area. In total, we genotyped about 1% of the genome, or 13 Mbp, of 149 individuals using RAD-sequencing (Table S5). Overall, we uncovered 11,489 single nucleotide polymorphisms (SNPs) in the 8 focal populations, and 17,288 SNPs if we included the scattered sample. Individuals were overall highly homozygous (>99%), indicating that self-fertilization dominates in Cologne populations. Populations showed a strong genetic structure, as can be seen from the genetic distance matrix (Figure 2a): Populations BER, GHU, RKO, and SHY consisted exclusively of genetically identical individuals, with the exception of a single plant with 99% heterozygosity from the GHU site. Populations BIS, HLU, KAS, and MIL each consisted of two genotypes. Overall Fst reached 0.72. Genetic similarity allowed clustering genotypes into three groups: Group 1 comprised RKO and BER, group 2 KAS-1, KAS-2, and BIS-2, group 3 consisted of HLU-1, HLU-2, and SGY. The genotype of plants collected in 2016 was the same as that of individuals collected in 2017 in the same location, confirming that local populations are most often formed from one (maximum two) single genotypes that persist over years.

**Fig. 2:**
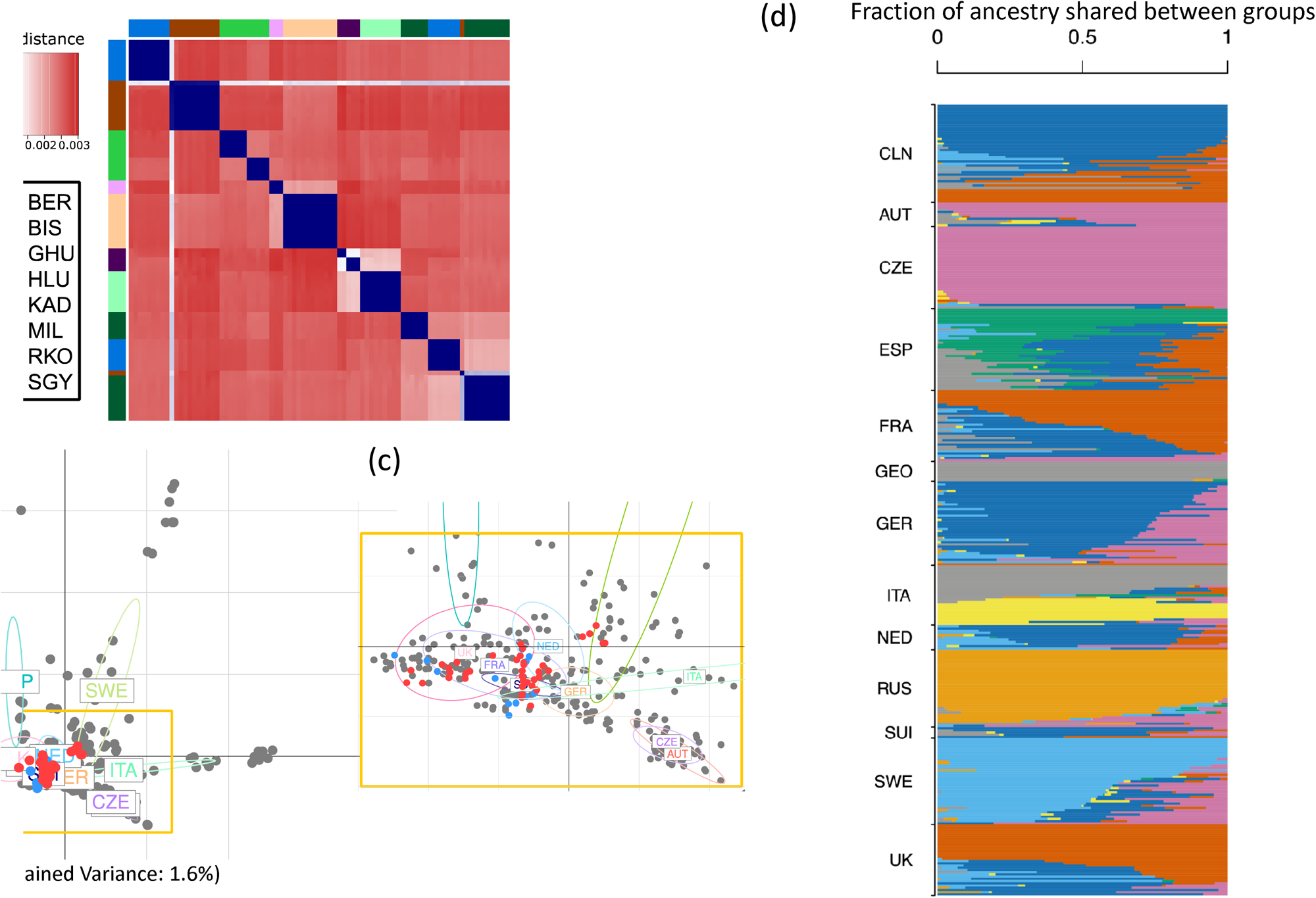
a) Analysis of genetic distance between progeny of 85 individual plants from 8 sampling sites in Cologne determined by RAD-sequencing. Individual plants are colour coded according to their origin. b) Principal component analysis of genetic differences between Cologne plants from the 8 sampling sites (blue), plants from additional sites of the sampling area (red) and 357 selected genotypes of the Arabidopsis 1001 genome set (grey). Origin of 1001 genome genotypes is indicated for countries represented by higher number of accessions (AUT-Austria, CZE-Czech Republic, ESP-Spain, FRA-France, GER-Germany, ITA-Italy, NED-The Netherlands, SUI-Switzerland, SWE-Sweden, UK-United Kingdom). c) Magnification of the yellow marked part of the plot b). d) Proportions of ancestry shared between Cologne genomes (CLN) and genomes collected in Europe was estimated by ADMIXTURE. On the basis of minimum cross-validation error, K=8 clusters were identified in the sample of 357 previously published European genomes. From top to bottom are shown genomes from Cologne, then from European dataset. The latter are grouped by country of origin and sorted by ancestry within groups.

In order to understand the origin of the genotypes that established populations in the city, we compared the Cologne samples to the European genotypes of the 1001-genome collection (Alonso-Blanco et al., 2016). We took a subset of representative genotypes from both datasets and first ran a principal component analysis (PCA) (see Materials and Methods). If habitats had been colonized via a stepping stone model, we would expect that the genotypes of the Cologne sample form a cluster of genotypes that stands out of the 1001 genomes. Cologne samples, however, co-localized with western and Central European accessions in the PCA, especially with accessions from Germany, France, the Netherlands, and the UK. They did not form a defined subgroup separate from the other genotypes (Figure 2b, c, Figure S6). Hierarchical clustering using admixture analysis confirmed that individual lineages share ancestry with individuals collected in Germany, the Netherlands or France in variable proportions (Figure 3d). In fact, the average number of pairwise differences between Cologne lineages was comparable to the number of differences between samples of the 1001 genomes (Figure S6). In conclusion, at each site, *A. thaliana* individuals formed either one or two clonal lineages, and collectively they do not form a local endemic group of similar genotypes. In addition, populations displayed no signal of isolation by geographic distance (Figure S6, Mantel test p>0.05) nor isolation by environmental distance (Mantel test p>0.05 for all environmental factors). We conclude that new populations are often established by long range migration and that the stepping stone model, by which new patches would be colonized by propagules of neighbouring patches, is inconsistent with the pattern of genetic diversity we observed (Duforet-Frebourg & Slatkin, 2016).

**Fig. 3:**
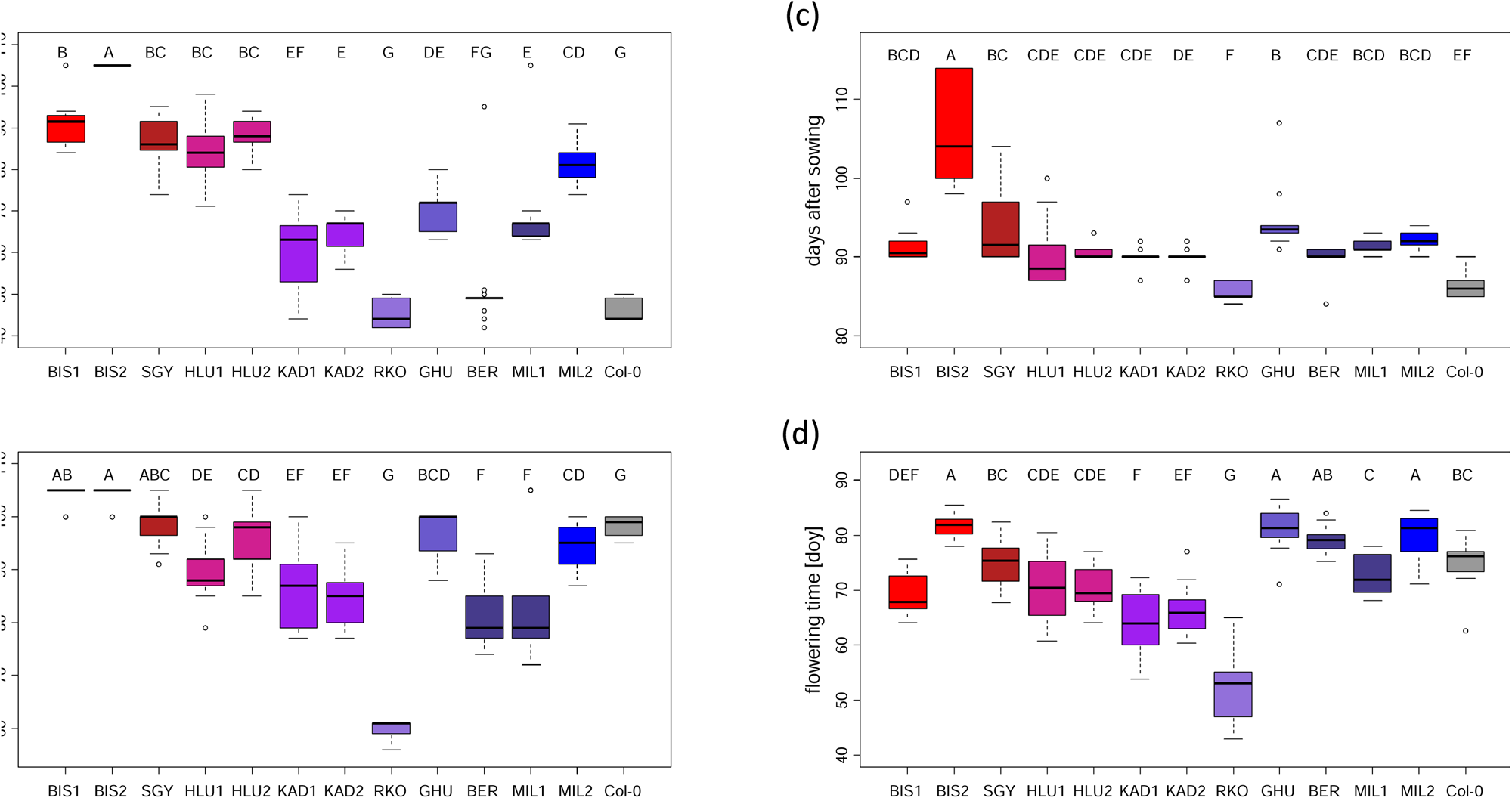
Flowering time (days after sowing) of plants grown in controlled conditions: long days (16 h light/20 °C; 8 h dark/18°C) (a), short days (10 h light/18 °C; 14 h dark/14 °C) (b), or short days followed by vernalisation (8 h light/4 °C; 16 h dark/4 °C) and shift to long days at day 64 (c). Flowering time (days of following year) of plants sown on dry soil on September 20^th^ and grown without artificial watering in the botanical garden (d). Letters indicate significant differences at a level p<0,05.

### Cologne urban populations show genetic variation in flowering time and dormancy regulation

Variation in genetic pathways regulating flowering time and germination may contribute to life-history differences recorded on-site. To link the germination and flowering behavior of the urban genotypes to knowledge about these well-studied pathways, we analyzed flowering under standard conditions in a randomized block design and included the well-studied genotype Col-0 as a reference. In total, three controlled environmental conditions were set: long days (LD), short days (SD), and a combination of short days/vernalization/long days (SDV). This setup thus allowed us to thoroughly examine genetic variation in the regulation of flowering time. The first two conditions allowed us to derive photoperiod-dependent differences in the regulation of flowering; the third mimicked the succession of environmental cues experienced in the fall, winter, and spring (Andrés & Coupland, 2012). Differences between LD and SDV conditions revealed variation in vernalization requirement, because it reveals the extent to which flowering time differences in LD tend to decrease in SDV as a result of exposure to the cold temperatures.

Flowering times differed between genotypes under all conditions (F_12,134to158_>39.53, p<2 e^-16^) and the interaction between genotype and experimental condition was signicant (F_24,461_= 36.72, p<2 e^-16^). Under LD conditions, BER and RKO plants flowered as early as the early flowering reference genotype Col-0, whereas BIS2 plants had not flowered after 105 days, indicating that the former do not require vernalization to flower (Figure 3a, Table S7). Under SD conditions, several genotypes flowered significantly earlier than Col-0, indicating that natural Cologne genotypes might flower in the short days of the fall or the late winter: RKO plants flowered earliest; BER, KAD1, KAD2, and MIL1 flowered only slightly earlier than Col-0. Both genotypes from the BIS site were unable to flower under SD conditions and were thus unlikely to flower in the fall (Figure 3b). When plants were grown under conditions simulating winter (SDV), most genotypes flowered within a narrow time period, with only BIS1 and BIS2 displaying significantly later flowering (Figure 3c). This experiment demonstrated that there is considerable genetic variation among the 12 Cologne genotypes for the regulation of flowering time. The magnitude of these differences depended on photoperiod and vernalization.

The genotypes also differed in how they regulated germination (Table S8). After three months of seed storage at room temperature, germination assays revealed that the genotypes differed in primary and secondary dormancy. When seeds were untreated, their germination rates indicated levels of primary dormancy (Baskin & Baskin, 2004; Finkelstein et al., 2008). Untreated seeds of BER, BIS2, MIL1, and MIL2 plants had high germination rates, whereas seeds of plants from the other genotypes had a less than 50% germination rate (χ^2^, df=11, N=48, p<2.2e^-16^; Figure 4a). Incubating non-dormant seeds at very low (-21°C) or high (35 °C) temperatures induced variable levels of secondary dormancy in Cologne genotypes, as indicated by a significant interaction between genotypes and germination conditions (likelihood ratio test, χ^2^= 83.2, df=24, p=1.81e^-8^, Figure 4c and supplemental Figure S7).

**Fig. 4:**
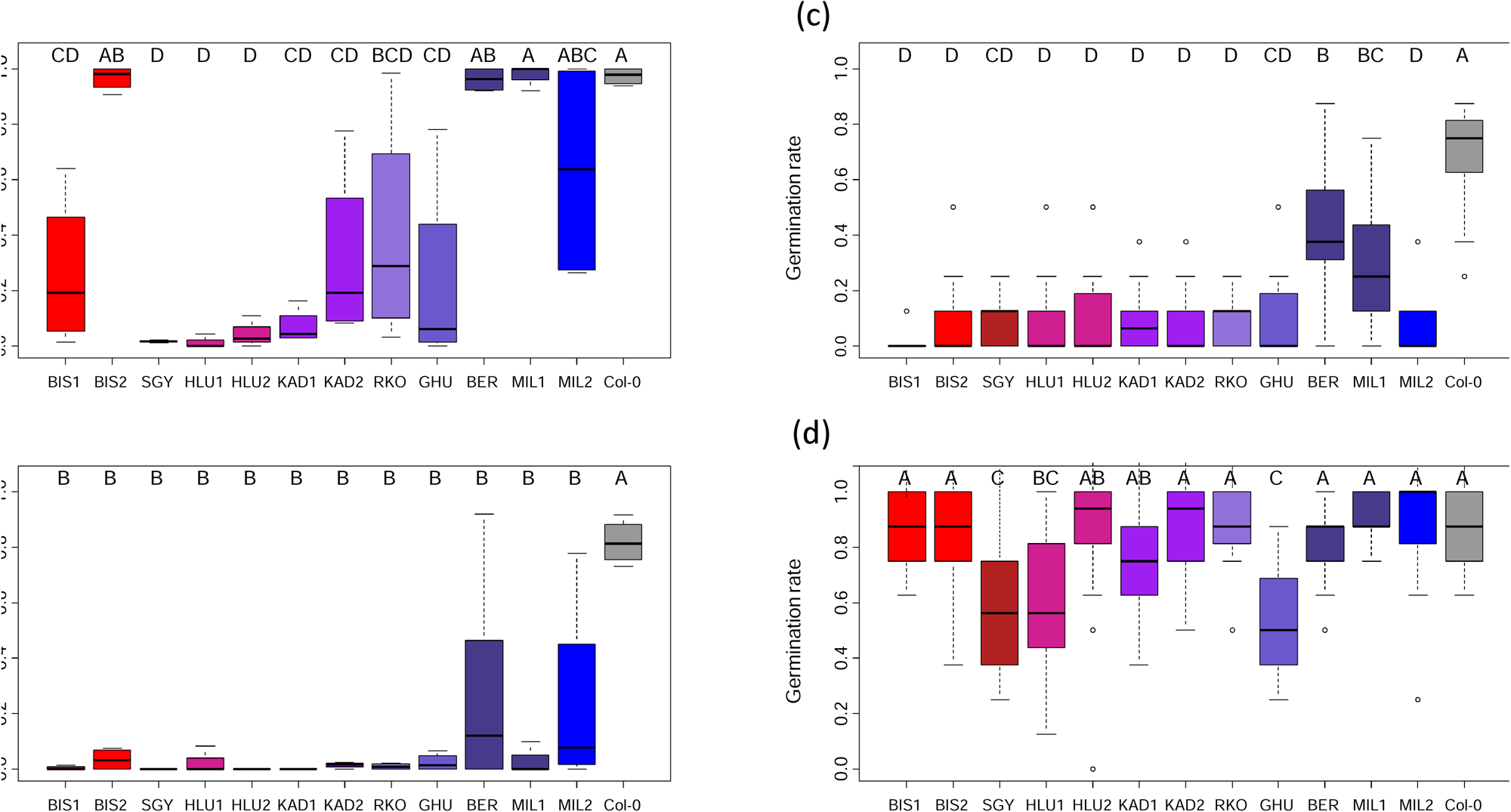
Germination frequency of seeds from the 12 different Cologne genotypes and Col-0 controls. Seeds were germinated on filter paper at 20 °C in petri dishes after 3-4 moths of afterripening to test for primary dormancy (a) or incubated at 35 °C in the dark for 4 days before incubation at 20 °C to observe heat induced secondary dormancy (b) (4 replicates per genotype). Germination frequency of seeds counted on October 5^th^, that were sown on dry soil in a common garden experiment without artificial watering on August 23^rd^ (c) or September 20^th^ (d) demonstrating different reaction to heat induced secondary dormancy under natural field conditions. Letters indicate significant differences at a level p<0,05.

### Outdoor common garden experiments confirm differences in germination and flowering regulation under field conditions

The 12 genotypes were further assayed in an outdoor common garden located in Cologne, and the genetic variation underlying their differences in survival and fertility was quantified. Here again, we included the widely used genotype Col-0 in the experiment as a reference (Table S9a-b). Since phenological variation is known to depend on the timing of germination, seeds were sown directly into pots at a field site in late August, mid-September, early November (6 weeks, 9 weeks, and 3 months after harvest), and the end of February. As pots were not watered artificially, germination and further development depended on natural rainfall and temperature. Germination rates among genotypes strongly differed in the August and September sowing cohorts. In the August cohort, 43 days after seeds were sown, only Col-0 plants had germinated to 100%; BER and MIL1 showed 43 and 28 % germination, respectively, 10 % or fewer of the seeds of the other genotypes had germinated (differences between the genotypes: F_12,195_=16.97, p<2e-16, Figure 4c). In the September cohort, 15 days after seeds were sown, most genotypes had germinated at 80% or more, and only HLU1, GHU, and SGY had germinated at less than 60% (after 22 d: F_12,195_=15.84, p<2e-16; Figure 4d). Eventually, most seed germinated, but the most delayed was the August cohort (Figure S8a-d). Later germination in this cohort can be explained by heat-induced secondary dormancy. In contrast, differences in germination for the November and February cohorts were minor (Figure S8e,f).

More than 95 % of the germinated seedlings survived until seed set. Most flowered in spring with the exception of the Col-0 and BER plants from the August cohort, which germinated almost immediately and flowered already in late autumn/winter. Differences in flowering time, which were observed for each of the four cohorts (F_12,195_>13.33, p<2 e^-16^), revealed a significant interaction between genotype and sowing date (F_34,755_= 20.89, p<2e^-16^, Table S3, Figure 3d, Figure S9). Flowering times observed for the 12 genotypes were generally not correlated across the various common garden cohorts grown outdoor, with the exception of the February and September cohorts (Figure S10). Fertility among the genotypes, determined by the number of fruits per pot, differed between the genotypes for the August, September, and November cohorts (F_12,195_=3.072, p=0.00054; F_12,194_=7.721, p=1.15e^-12^; and F_12,195_=4.702, p=1.01e^-06^). Again, a significant interaction between genotype and seed-sowing date was observed (F_24,584_=3.534, p= 4.75e^-08^). For the August cohort, only two genotypes (RKO and SGY) were less fertile than most of the genotypes (Table S3, Figure S11). Fitness differences for the plants grown from seeds sown in February, all of which germinated within a week in early March and flowered 23-39 days later, were not significant. With 52% of the plants dying after an unusually warm dry spell in May, the average fitness of the spring cohort was low.

In summary, plants from the 12 Cologne genotypes displayed remarkable genetic differences in their ability to control germination and flowering in common garden experiments. Flowering times measured in controlled growth chamber and in the common garden were significantly correlated, but correlation depended on the various experimental indoor conditions and/or the seed dispersal times characterizing the different common garden cohorts (Figure S10).

### Environmental variation among sites of origin correlates with genetic variation observed in common gardens and controlled environments

The phenotypes of urban *A. thaliana* individuals in situ correlate with the ecological characteristics of the sites in which they are monitored. We further tested whether the distribution of genetic differences revealed in common gardens and under lab conditions is independent of the ecological characteristics at the sites of origin. We found that genetic variation in the rate of germination under late summer conditions in the August cohort or after cold treatment correlates positively with variation in the severity of disturbance (r=0.763, p=0.0039 and r=0.816, p=0.0012, respectively). Furthermore, in the absence of vernalization, variation in the time to flowering correlates negatively with variation in disturbance severity and temperatures (e.g., indicator value T (flowering date in LD or SD): r=-0.814 and -0.806; p=0.0013 and 0.0016; Figure 5, Figure S12). Genetic variation in phenology is thus not randomly distributed across habitats. Its distribution matches the disturbance gradient along which the *A. thaliana* plant populations monitored in this study are distributed.

**Fig. 5:**
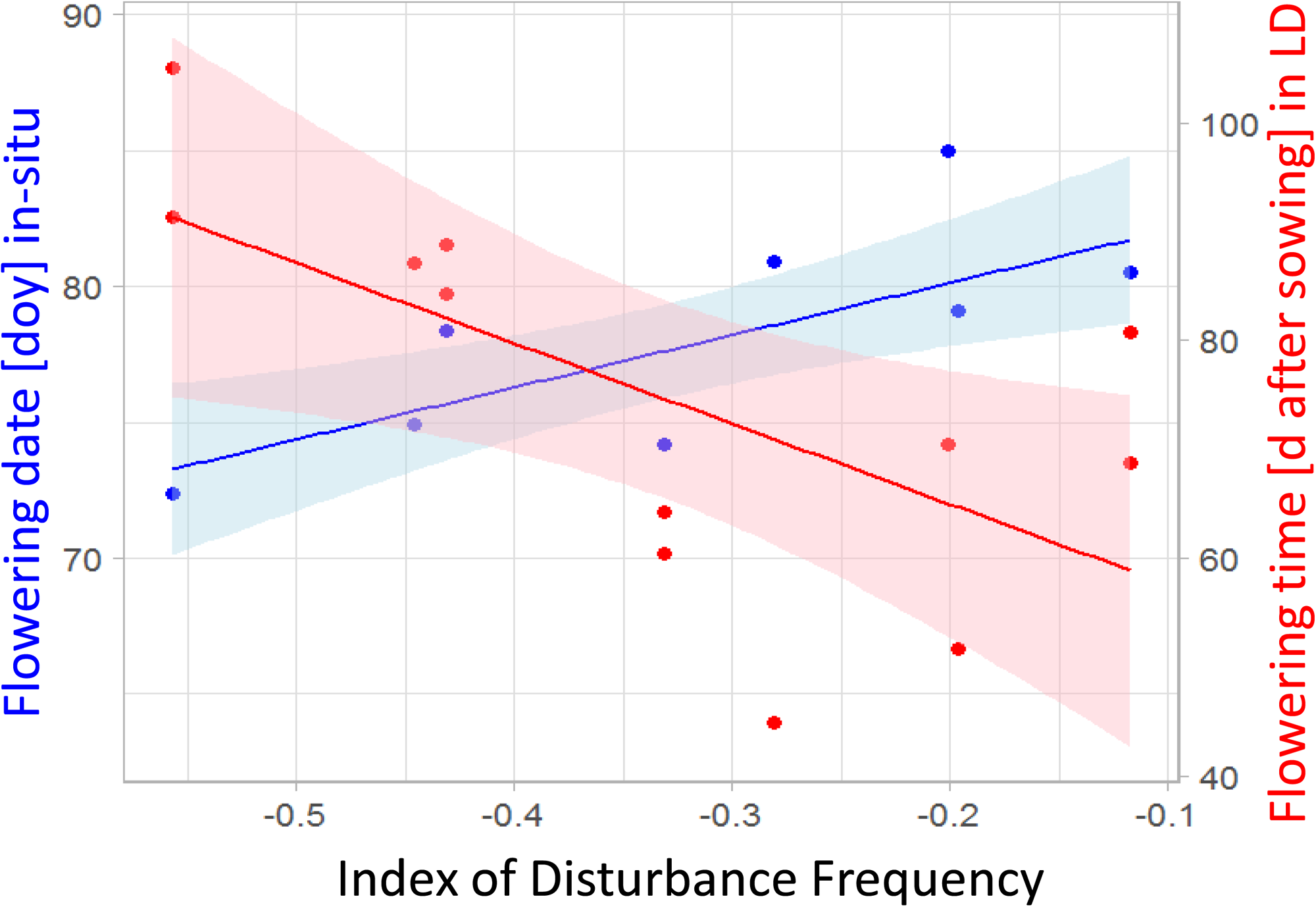
Variation of phenology in urban *A. thaliana* genotypes, as observed within urban habitat patches in the City of Cologne (in-situ; blue), and in a common-garden experiment in the same city (ex-situ; red) as a function of variation in the index of Disturbance frequency at the site of origin. The timing of flowering in-situ (blue) increases with disturbance frequency, whereas the opposite trend is observable for the timing of flowering in indoor common garden conditions (red).

Both genetic variation for the regulation of flowering time in LD conditions and phenotypic variation for flowering time *in situ* were correlated with disturbance frequency, but the slope of the relationship changed signed (F_1,20_=14.6984, p=0.001, Figure 5, Figure S13). We conclude that genetic variants accelerating flowering in the absence of vernalization are found in the most disturbed sites, where plants are observed to flower at a later date.

## Discussion

Urban habitats form a mosaic of diverse environments that challenge plant growth in different ways. However, the impact of this heterogeneity on the distribution of adaptive genetic variation is not known (Rifkin, 2018). While the influence of urban heterogeneity on gene flow and population differentiation has been reported in animal and plant species, its role in shaping the distribution of genetic variation for adaptive traits remains unexplored (Beninde et al., 2015; Gorton et al., 2018; Rivkin et al., 2019). By combining plant community and trait-based ecological measurements with an in-depth exploration of genetic diversity at the genomic and phenotypic levels, this study bridges this knowledge gap. The results provide compelling evidence that the genetic variation in *A. thaliana* is not randomly distributed across environmentally diverse urban habitats. These findings enhance our understanding of the actual ecology of a species that serves as a model in plant molecular biology (Takou et al., 2019), and further provide valuable insights into adaptive genetic traits in urban environments (Rivkin et al., 2019).

*A. thaliana* populations sampled here confirm previous reports about urban populations (Bomblies et al., 2010): Each habitat patch is colonized by one, sometimes two, clonal lineages. An analysis of genomic variation showed that these lineages belong to the broadly distributed gene pool of northwestern European *A. thaliana* genotypes, supporting the notion that urban city patches were established from a pool of regionally diverse migrants. Furthermore, our study shows that the urban habitats hosting spontaneously occurring *A. thaliana* stands are heterogeneous. The advantage of ecological characteristics based on community indicators, which leverage the particularly extensive information available for the German flora, is that they provide robust measures of environmental variation (Diekmann, 2003; Fanelli et al., 2006). Here, these characteristics reveal that *A. thaliana* habitats form a continuum of varying levels of humidity and nutrients, as well as varying frequencies of disturbance. The ecological differences we quantified were clearly associated with the phenotypic differences displayed by plants *in situ,* as plants growing under better edaphic conditions (more nutrients, more water) were larger, and plants growing in disturbed habitats bolted later and bore more fruits. *In situ* observations of plant diversity, however, do not allow disentangling the effects of plant plasticity to the environment from the effect of genetic differences between populations (Aguilar-Trigueros et al., 2017; Cadotte & Tucker, 2017; Thakur & Wright, 2017). Unraveling whether the successful occupation of diverse urban habitat patches is solely facilitated by plant plasticity represents a critical step towards understanding the ecological dynamics of biodiversity in urban environments (Brans et al., 2017; Rivkin et al., 2019).

Focusing on the 12 clonal lineages identified by genomic analysis, we thus explored whether genetic variation contributes to the phenological variation observed *in situ*. By comparing the lineages forming each population patch under common growth conditions, we found strong genetic differences in how they regulate seed germination and flowering. In total, we relied on at least seven indoor and outdoor common garden conditions to show that the 12 clonal lineages varied in their requirements for vernalization and their ability to flower during shortening daylength, as well as in the ability of their seeds to delay germination after exposure to cold and heat. We confirm previous findings that the timing of germination plays a major role on life-history traits manifested later in the life cycle (Donohue, 2009; Wilczek et al., 2009). By contrasting variation in plant traits expressed *in situ* with *ex situ* observations made in common garden settings, we showed that genetic variation in phenology tends to buffer the effect that environmental heterogeneity has on trait variation. Indeed, we found that genotypes that flowered early under LD or SD conditions (*ex situ*) were found in sites that were more disturbed. Whereas, those from the less disturbed sites flowered later under LD or SD conditions but flowered early *in situ* (Figure 5). Indeed, the timing of flowering in LD conditions and *in situ* showed opposite patterns of covariation with disturbance frequency (Figure 5). Therefore, genotypes growing in habitats with relatively harsher environmental conditions (characterized by resource stress, frequent disturbance, and a delay in development) tend to have traits genetically tailored to a life cycle that is shorter in the absence of vernalization. We thereby reveal a remarkable pattern of counter-gradient adaptation (Conover & Schultz, 1995), wherein genetic variation for flowering time in LD reduces the extent of phenological variation that would otherwise typify the heterogeneous environments hosting *A. thaliana* in the city. Therefore, variable genetic traits may not always be directly observable across environmental gradients. Counter-gradient variation is indeed a well-known confounding experimental error for *in-situ* studies, that can only be resolved with genetic studies in common gardens (Conover & Schultz, 1995).

We believe that, in this study, the distribution of genetic differences in the regulation of flowering reflects the outcome of a process driven by local selective pressures. Indeed, we found no correlation between genetic distance and geographic or ecological distance, so the non-random pattern of co-variation between disturbance level and genetic variation in life-history cannot be explained by local seed dispersal. We identify two other potential explanations: i) specialized genotypes have adapted via the selection of new mutations affecting life history in an otherwise non-adapted lineage (Orr, 2005), or ii) local environments filtered mis-adapted genotypes out, in the same way that they exclude mis-adapted species (Guo et al., 2018; Kraft et al., 2015; Laliberté et al., 2014; Yang et al., 2022). Since many alleles altering the genetics of flowering time have already been characterized in the western European region from which the populations originate (Le Corre, 2005; Lopez-Arboleda et al., 2021), it is highly likely that genotypes with diverse regulatory alleles were among the pool of migrants that populated habitat patches in the city of Cologne. Environmental filtering thus appears to be the most parsimonious explanation for the non-random distribution of genetic variation in flowering time documented in this study. Considering that the assessment of variation *in situ* and at the genetic level constrained us to the study of only 8 populations, our findings suggest that urban environmental heterogeneity has a strong filtering power on the genetic basis of intraspecific trait variation. Environmental filters imposed by habitat heterogenetiy in cities have been shown to diversify plant communities (Aronson et al., 2016). Indeed, cities can preserve species diversity, depending, for example, on the size and connectivity of habitats within cities (Beninde et al., 2015; Hahs et al., 2009). To the best of our knowledge, this study is the first to show that environmental filtering also has the potential to maintain functionally diverse and locally adapted genetic lineages. Such lineages form *bona fide* ecotypes, a term often used in the literature but seldom scrutinized.

Our assessment of *in situ* and *ex situ* variation in phenology highlights the importance of interdisciplinary approaches in ecology, quantitative genetics, and genomics. Indeed, our study identifies a pattern of adaptive genetic trait variation that is not directly observable in situ. With our sample of a small number of highly structured populations composed of one or two clonal lineages, we do not yet have the power to identify the loci responsible for adaptive life history variation (Hancock et al., 2011; Herden et al., 2019). The number of variants that contribute to the genetic variation of plant phenology in Cologne *A. thaliana* populations will have to be determined in future studies.

Finally, one can speculate that disturbances may be the driving force behind the environmental filtering of genetic variation in life history within the city. However, in urban plant communities, environmental filters are shaped by a complex interplay of various factors, including urban habitat transformation, fragmentation, typical urban alterations such as heat islands or pollution, and human preferences(Williams et al., 2009). It is highly likely that each of these factors co-varies with the degree of disturbance. Consequently, future studies focusing on genetic variation within cities should intend to disentangle the specific factors filtering genetic variation in life history traits within urban plant populations and communities.

## Acknowledgments

We thank Denitsa Ilieva for help in common garden experiments, and Emily Wheeler, Harwich, MA, for editorial assistance. We acknowledge funding from the German Science Foundation in the framework of TRR341 “Plant Ecological Genetics” and from the Center of Excellence in Plant Sciences (CEPLAS), in Cologne and Düsseldorf.

## Authors’ Contributions

AL, JM, GS, AS conceived the ideas and designed the methodology; JM, GS and AL wrote the manuscript; AF, GS, HD, JT, GC collected and/or analyzed the data; all authors contributed to the draft and gave final approval for publication. Sequencing data was deposited as submission numbers ERS5063293 to ERS5063441, project PRJEB40091 in the ENA repository. We acknowledge funding from the Deutsche Forschungsgemeinschaft within the framework of TRR341.

