## Supplementary text and figures for "Environmental filtering of life-history trait diversity in urban populations of *Arabidopsis thaliana*"

### Journal of Ecology Supporting Information

**Environmental filtering allows local adaptation of life-history traits of *Arabidopsis thaliana* urban populations**

**Supplementary information:**

**Bioinformatics pipeline and identification of A. thaliana population structure in the city of Cologne**

Genomic DNA sequence was characterized by RAD-Seq as described by Dittberner at al. (2019) using 10 pools with 20 plants each. The mapping of sequenced reads and the determination of population structure is described in the following.

Reads were trimmed with TRIMMOMATIC and low quality reads were sorted out using FastQC. Reads were subsequently mapped to the *A. thaliana* reference genome TAIR10 (Berardini et al., 2015). Minimum coverage for SNP calling was 15. Forty-five individuals with missing data in more than 70% of loci were removed from the main dataset (see below). Loci with more than 20% missing data were discarded. The RAD genotype dataset was merged with the 1001 genomes SNP dataset (Alonso-Blanco *et al.* 2016, Table S11) using *bcftools* (Li, 2011) and filtered to include only loci genotyped in the RADseq dataset. Genetic distances between pairs of individuals were calculated using a custom python script. A heatmap of genetic distances was created using the *heatmap3* package (https://CRAN.R-project.org/package=heatmap3). Genetic distance among populations was calculated using the *hierfstat* package (Goudet, 2005). For each environmental factor, pairwise Euclidean distance among populations was calculated. Correlation between environmental and genetic distance as well as between geographic and genetic distance was tested using a Mantel test with 999 permutations.

To relate Cologne genotypes to the larger collection of accessions of the 1001-genome project, we included accessions from Germany and from countries close to Germany in western, northern or southern direction (France, Spain, Switzerland, Netherlands, United Kingdom and Italy) with at most 40 accessions per country. Accessions from countries with more than 40 accessions in the dataset were randomly subsampled and relict accessions (Alonso-Blanco et al., 2016) were removed. Cologne genotypes from both sample sets (‘populations from the eight habitat patches’ and from ‘scattered sites’) were subsampled to remove genetically identical individuals, resulting in a set of 51 genotypes. In order to visualize similarity between plant genomes, a principal component analysis was conducted using the *vcfR* package (Knaus & Grünwald, 2017) and the *adegenet* package (Jombart & Ahmed, 2011) with missing variants scaled to the mean.

To assign genotypes to the 45 individuals with low coverage, we called SNPs again allowing for minimum coverage of 2. We removed all SNPs with more than 50% missing data. We used the software *SNPmatch* (Pisupati et al., 2017) to create a database of the high quality SNP dataset and find the closest match to the database for each of the 45 low-coverage individuals. If the 3 best matches were the same genotype and identity was over 97%, we assigned that genotype to the individual.

We determined the origin of the 52 main lineages sampled in Cologne by comparing their diversity to a representative subset of 415 lines included in the 1001 genomes (Table S11). We first ran a Principal Component Analysis (PCA). FWe analyzed hierarchical population structure with

ADMIXTURE v1.3 (Alexander et al., 2009). To this end, we filtered out lines that had more than 70% missing SNP information and removed SNPs that were present in only one individual. Then, to eliminate samples that share a recent common ancestry, we computed the density of pairwise differences for every pair of genomes, and discarded one genome for every pair with less than 0.01 differences., reducing the dataset to 355 lines and the number of SNPs from 46,457 to 29,593. In total, we included in the admixture analysis 44 Cologne genomes (CLN), 32 genomes from France (FRA), 37 from Czechia (CZ), 11 from Austria (AUT), 39 from Sweden (SWE), 32 from the UK, 38 from Germany (GER), 35 from Russia (RUS), 11 from the Netherlands (NED), 37 from Spain (ESP), 5 from Switzerland (SUI), 27 from Italy (ITA) and 9 from Georgia (GEO). The dataset was then pruned in in PLINK v1.90b3.40 for linkage < --indep-pairwise 50 10 0.1 > and for missingness < --geno 0.1 > (Chang et al., 2015). We ran Admixture with k varying from one to thirty, and chose as the best fit the run with the lowest cross-validation error. The optimal number of clusters is the number that has the lowest cross-validation error (cve). In this data optimal k was 8, a number roughly similar to what was previously reported (Alonso-Blanco et al., 2016). Genome ancestry was shown starting from the Cologne data set and followed by 1001 genomes sorted by country of origin in alphabetical order. Within each group, genomes were sorted by ancestry proportions at the cluster (color) that was most represented on average among lines of each country.

#### SUPPORTING TABLE LEGENDS

**Table S1:** Phenological stages and fitness proxies measured *in situ* for urban populations of *A. thaliana*; with units of measurement; minimum, maximum (Min/Max) and median trait values for the whole data collected across the study sites. doy: days of year, mm: millimetre.

**Table S2:** Flowering dates and fertility of Cologne lines and Col-0 grown in common garden experiments (averages and standard deviation). Seeds were sown on dry soil on August 23, September 20, November 8, or February 6, and grown without artificial watering. Letters indicate significant differences at a level p<0,05.

**Table S3:** GPS coordinates of the sites monitored in the study

**Table S4:** Plant community present in the sites of the study and associated Ellenberg and disturbance indicator values (EIV and DIV, respectively).

**Table S5:** Summary of RAD-sequencing data. First generation progeny of plants described below in Table S6 were used for DNA isolation. Links to sequencing data are given for plants with high sequencing coverage. Assignements of individual plants to genotypes is described in material and methods.

**Table S6:** Phenology data of individual plants from the 8 monitored sites of Table S3. Each plant has a unique identifier containing the name of the sub-site of sampling.

**Table S7:** Flowering time data for individual plants grown in controlled conditions (indoor common garden). Data show the days when the first flower has open petals and is calculated from the day of sowing. NA- plants were lost during the experiment. Plants that did not flower within 105 days were assigned “>105”.

**Table S8:** Germination in controlled conditions (indoor common garden). Data indicate the fraction of germinated seeds after 10 d incubation at 20°C. Pre-treatments are described in material and methods.

**Table S9:** a- Summary of common garden experiments for plants sown in August, September and November and b- in February (see methods). Germination was recorded at different time points after sowing. For the February cohort, density was reduced to 1 plant per pot, for the other cohorts all plants were left to grow. For these experiments, flowering time is the average of flowering time of surviving plants.

#### SUPPORTING FIGURE LEGENDS

**Figure S1:** Characteristics of eight urban *Arabidopsis thaliana* populations and their habitat conditions in the city of Cologne, with (a) differences between habitat patches based on plant species' Ellenberg indicator for humidity (F), (b) light intensity (L) index , and (c) Herb layer structure-based disturbance index. Letters indicate significant differences at p<0.05.

**Figure S2:** Phenological stages and plant functional traits of the eight urban *A. thaliana* populations measured in-situ, with a) differences in bolting ($F_{7,149}$=17.81, p<$2e^{-16}$); b) differences in maximum rosette diameter ($F_{7,163}$=10.92, p=$2.85e^{-11}$). Letters indicate significant differences at p<0.05. c) correlations of populations' average phenological stages and functional traits with abiotic habitat characteristics given by the accompanying plant community (L -light, T -temperature, K -continentality, F -humidity, R -pH, N –nitrogen content) and disturbance regimes (disturbance frequency DF, severity DS, herb-layer structure DV).

**Figure S3:** Significant correlations between Ellenberg Indicator Values (EIVs) and Disturbance Indicator Values (DIVs) for the eight *A. thaliana* habitat patches in the City of Cologne. EIVS are light regime (L), temperature (T), continentality of climate (K), soil moisture (F), soil reaction (R), and nutrient availability (N); DIVs are Disturbance frequency index (DF), Disturbance strength index (DS), and Herb layer structure-based disturbance index (DV)

**Figure S4:** Development of phenological stages over time observed at the eight Cologne populations shown as percentage of plants reaching the stage of bolting, onset of flowering and onset of fruiting.

**Figure S5:** Correlation of average phenological stages and growth phenotypes between urban populations (only significant correlations are shown).

**Figure S6: a-**Pairwise genetic distance of genotypes sampled in the eight urban habitat patches (BER, BIS, GHU, HLU, KAD, MIL, SGY, RKO) and from additional scattered sites of the study area.

Genetic distance between genotypes, as measured by the average number of pairwise nucleotide differences p, ranged from 0.0017 to 0.0034 with a mean of 0.003. Genetic similarity allowed clustering genotypes into three groups: Group 1 comprised RKO and BER, group 2 KAS-1, KAS-2, and BIS-2, group 3 consisted of HLU-1, HLU-2, and SGY. Populations displayed no signal of isolation by geographic distance (Mantel test p>0.05) or isolation by environmental distance (Mantel test p>0.05 for all environmental factors). In the additional populations, 44 individual genotypes were found among the 65 sequenced, and no genotype was found at more than one site. Five accessions (RUS16, RUS17, ROD8, RUS23, RUW1) deviated from the rest and co-localized with Swedish accessions. Interestingly, we also found four individuals originating from a single site (ROD-1, ROD-2, ROD-3, ROD-4), which were virtually identical (1 SNP) to the accession TueWa1-2 from Tübingen (Germany), which was described in the Arabidopsis 1001 Genomes Project. This lineage might result from long-range migration.

b- Distribution of genetic distance between genotypes, as measured by the average number of pairwise nucleotide differences p, ranged from 0.0017 to 0.0034 with a mean of 0.003. Left panel shows the origin of the lines used in the study. Lines collected in Cologne (CLN) are practically as different from each other as lines from the broader sample including lines from neighbouring countries. This suggests that Cologne population were established from a diverse pool of migrant genotypes, presumably via long range migration.

**Figure S7:** Germination frequency of seeds from the 12 different Cologne genotypes and Col-0 controls. Seeds were germinated on filter paper at 20 °C in petri dishes following stratification for 7d at 4 °C and demonstrating that seeds are viable (a) or an incubation at -21 °C in the dark (b). (4 replicates per genotype).

**Figure S8:** Germination frequency of seeds from the 12 different Cologne genotypes and Col-0 controls germinated in common garden conditions. Seeds were sown on dry soil on August 23rd (a, b), September 20th (c, d), November 8th (e), or February 6th (f). Germination was scored on October 5th (a, c), October 31st (b, d) for the August and September sowing. Note that at the second time point, more than 50% of seeds germinated, but GHU, SGY, KAS1, and RKO still had approximately 20% lower germination rates compared to the other genotypes. Germination was scored on December 13th (e), and April 5th (f) for the November and February cohorts, respectively. Differences between a) and c) can be explained by high temperature induced secondary dormancy.

**Figure S9:** Flowering time (days of year) of plants from the 12 different Cologne genotypes and Col-0 controls grown in common garden conditions. Seeds were sown on dry soil on February 6th (a), August 23rd (b), September 20th (c), or November 8th (d). Letters indicate significant differences at a level p<0,05.

**Figure S10:** Correlation among averages for flowering time phenotypes measured in controlled laboratory conditions and in common garden experiments. Flowering times between the 4 planting cohorts show little correlation

**Figure S11:** Common garden experiments - number of siliques of single plants sown in February (a), or number of siliques per pot with 8 seeds sown in August (b), September (c), or November (d). Letters indicate significant differences at a level p<0,05.Scales differ for the plots.

**Figure S12:** Correlation of averages of phenotypes of plants in controlled lab conditions, outdoor common garden experiments (cohorts) and averages of Ellenberg and disturbance indicator values of the sites of their origin (L- Light, T- Temperature, K- Continentality, R- pH, N- nutrients, DF- disturbance frequency, DS- disturbance severity, DV- structure-based disturbance index). Germination rates were monitored at several time points after sowing. Flowering time was scored in controlled growth chamber under long days, short days, and in a sequence combining short days, vernalization and then long days (vernalization conditions). Germination was scored in controlled conditions on untreated 2-month-old seeds and after stratification. Secondary dormancy was determined by scoring germination in stratified seeds exposed to either freezing conditions or heat (see methods for details). Fertility was measured in all four outdoor common garden cohorts, but not in growth chambers.

**Figure S13:** Correlation of averages of phenotypes of plants in their original sites, in controlled lab conditions and in common garden experiments. Germination rates were monitored at several time points after sowing. Flowering time was scored in controlled growth chamber under long days, short days, and in a sequence combining short days, vernalization and then long days (vernalization conditions). Germination was scored in controlled conditions on untreated 2-month-old seeds and after stratification. Secondary dormancy was determined by scoring germination in stratified seeds exposed to either freezing conditions or heat (see methods for details). Fertility was measured in all four outdoor common garden cohorts, but not in growth chambers.

Journal of Ecology Supporting Information

### Environmental filtering allows local adaptation of life-history traits of *Arabidopsis thaliana* urban populations

**Authors:** Gregor Schmitz, Anja Linstädter
, Anke S. K. Frank, Hannes Dittberner, Andrea Schrader, Karl-Heinz Linne von Berg, George Coupland, Juliette de Meaux

#### SUPPORTING FIGURES

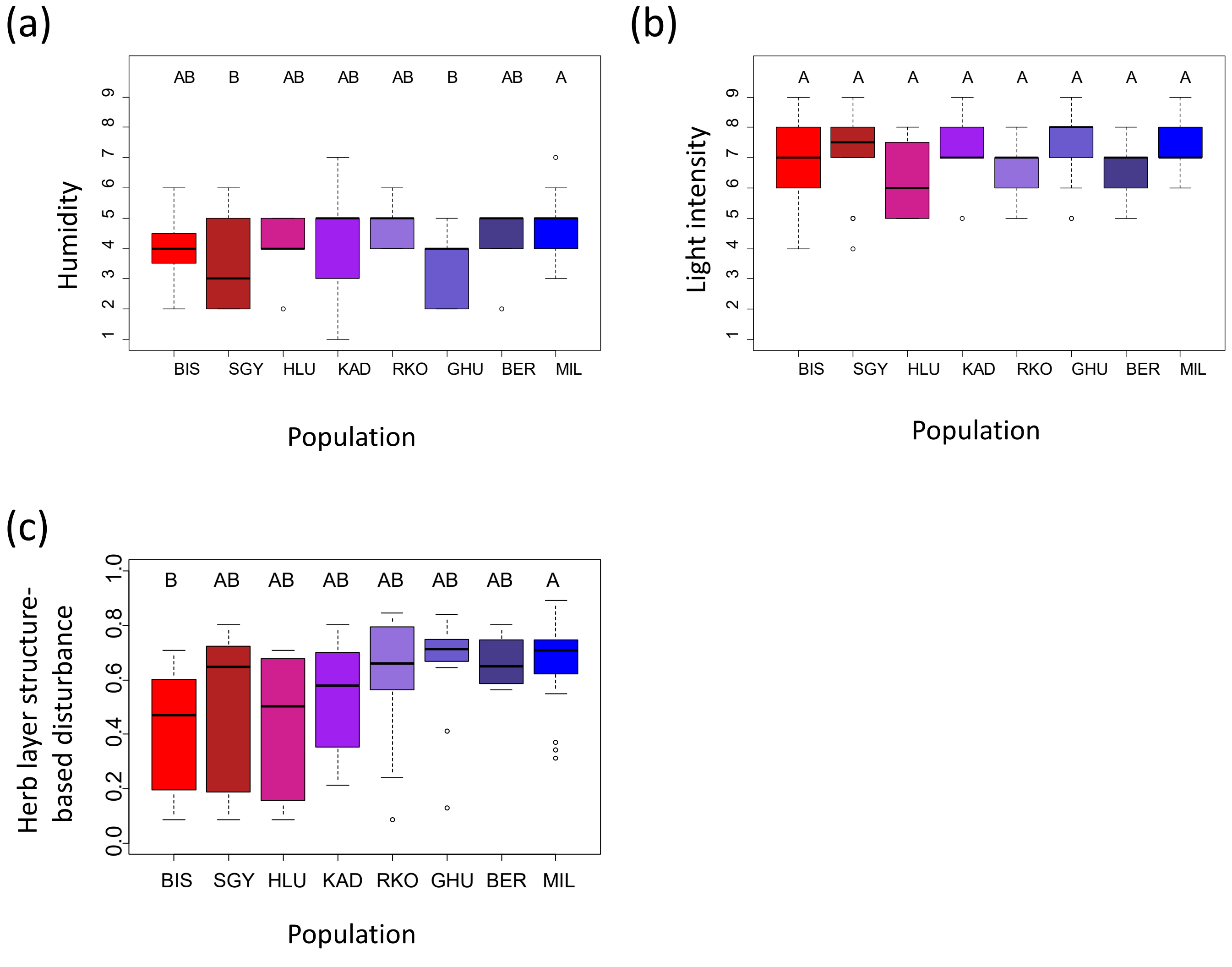


**Figure S1:** Characteristics of eight urban Arabidopsis thaliana populations and their habitat conditions in the city of Cologne, with (a) differences between habitat patches based on plant species' Ellenberg indicator for humidity (F), (b) light intensity (L) index , and (c) Herb layer structure-based disturbance index. Letters indicate significant differences at $p<0.05$.

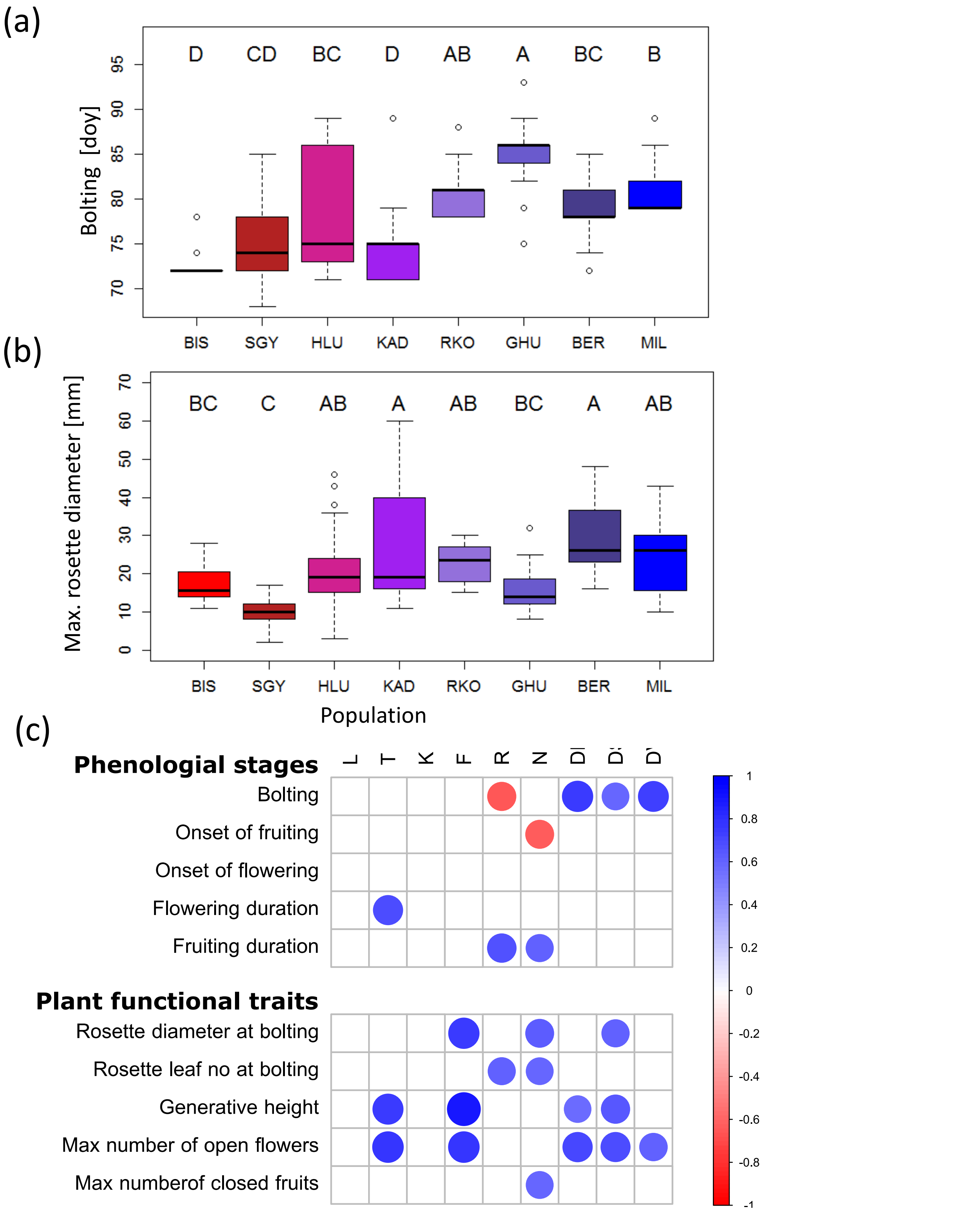


**Figure S2:** Phenological stages and plant functional traits of the eight urban *A. thaliana* populations measured in-situ, with a) differences in bolting ($F_{7,149}$=17.81, p<2e$^{-16}$); b) differences in maximum rosette diameter ($F_{7,163}$=10.92, p=2.85e$^{-11}$). Letters indicate significant differences at p<0.05. c) correlations of populations' average phenological stages and functional traits with abiotic habitat characteristics given by the accompanying plant community (L -light, T -temperature, K -continentality, F -humidity, R -pH, N –nitrogen content) and disturbance regimes (disturbance frequency DF, severity DS, herb-layer structure DV).

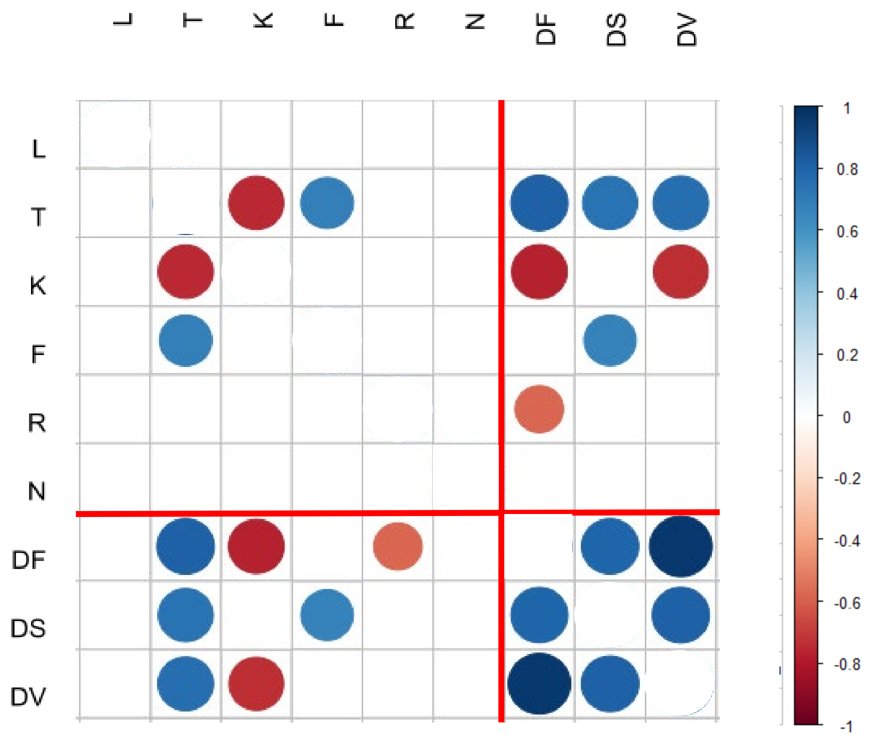


**Figure S3:** Significant correlations between Ellenberg Indicator Values (EIVs) and Disturbance Indicator Values (DIVs) for the eight *A. thaliana* habitat patches in the City of Cologne. EIVS are light regime (L), temperature (T), continentality of climate (K), soil moisture (F), soil reaction (R), and nutrient availability (N); DIVs are Disturbance frequency index (DF), Disturbance strength index (DS), and Herb layer structure-based disturbance index (DV)

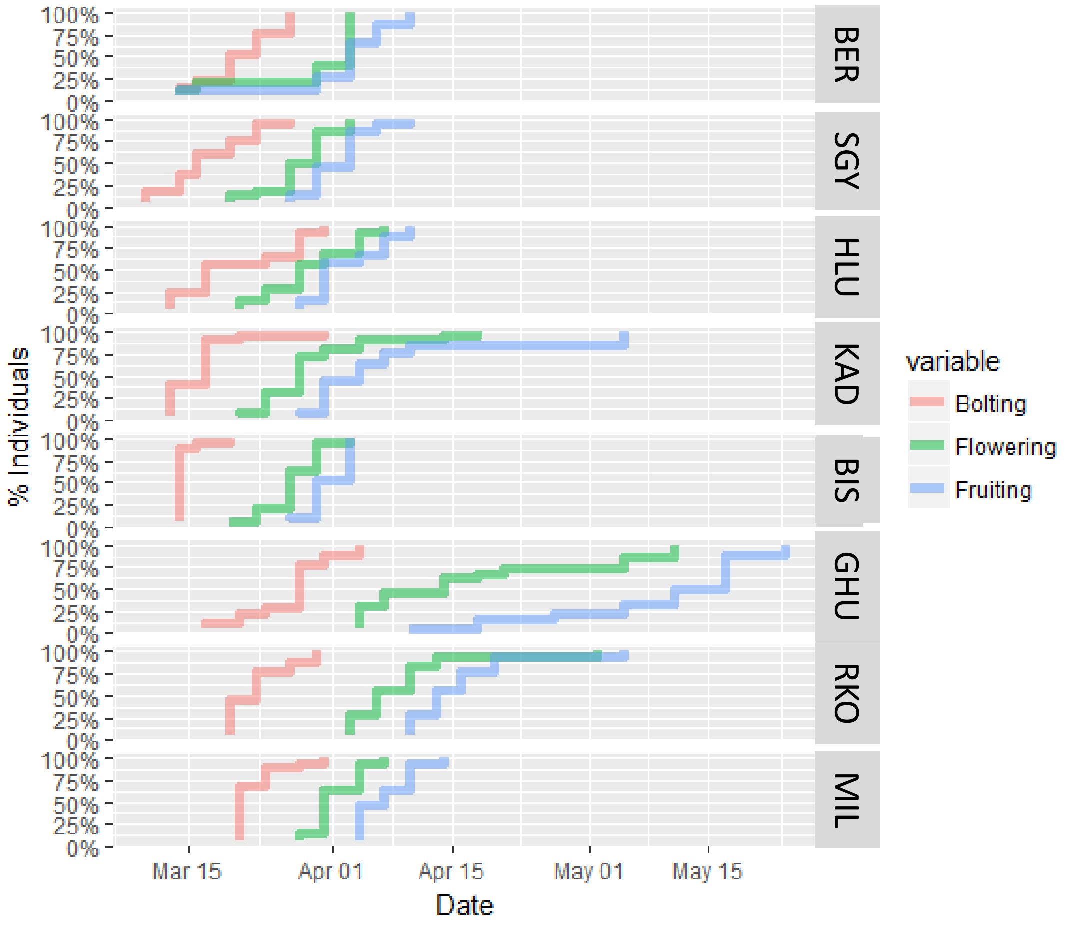


**Figure S4:** Development of phenological stages over time observed at the eight Cologne populations shown as percentage of plants reaching the stage of bolting, onset of flowering and onset of fruiting .

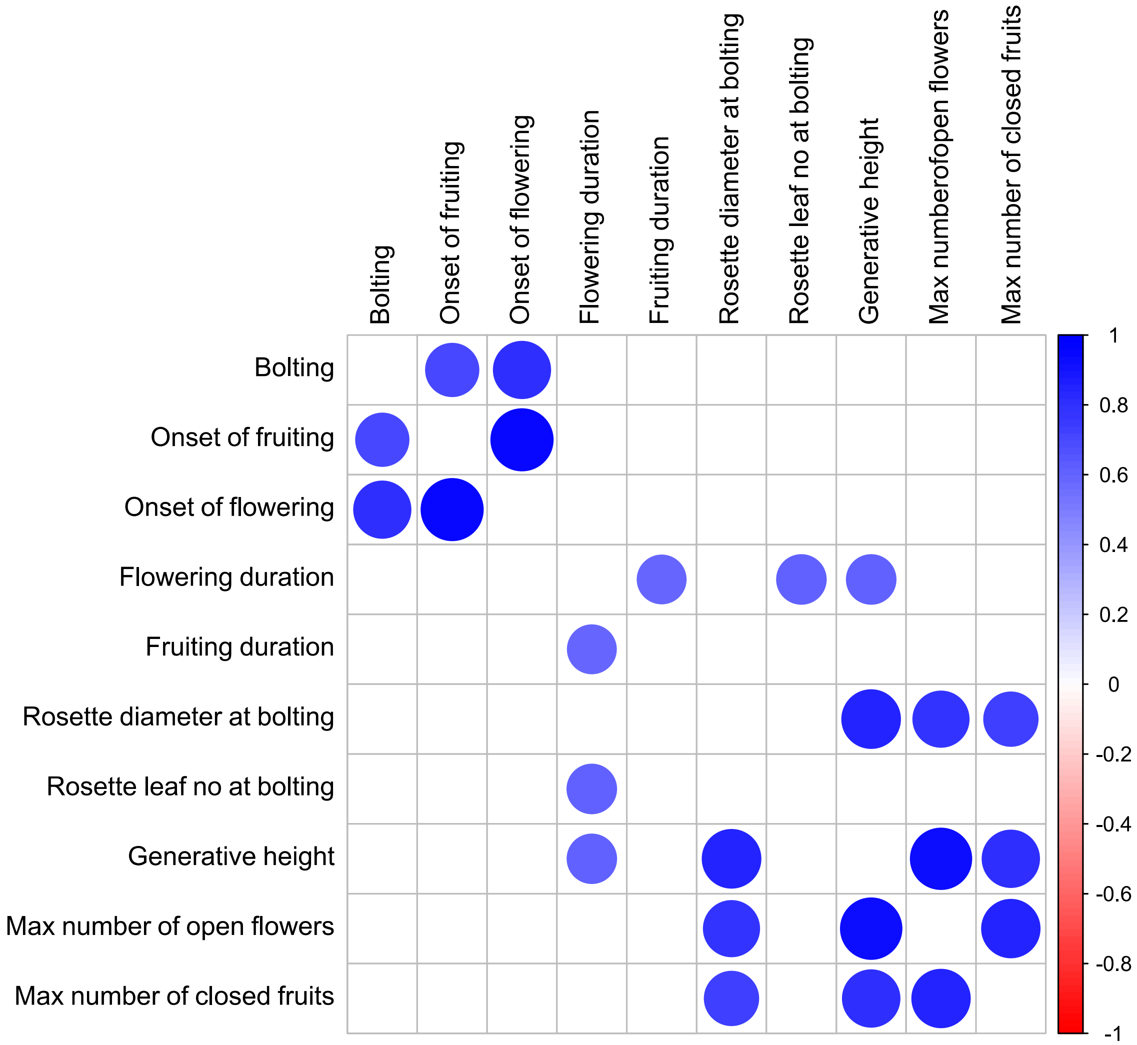


**Figure S5:** Correlation of average phenological stages and growth phenotypes between urban populations (only significant correlations are shown).

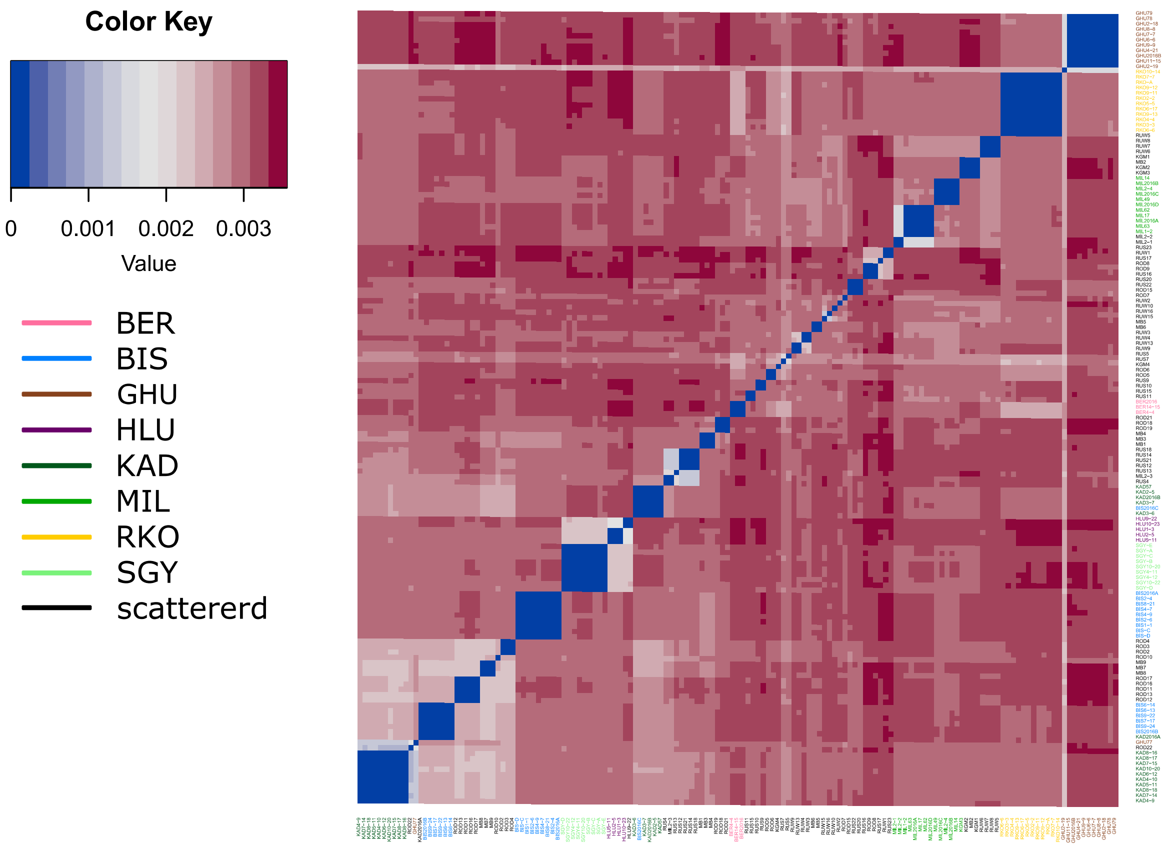


**Figure S6A:** Pairwise genetic distance of genotypes sampled in the eight urban habitat patches (BER, BIS, GHU, HLU, KAD, MIL, SGY, RKO) and from additional scattered sites of the study area.

Genetic distance between genotypes, as measured by the average number of pairwise nucleotide differences p, ranged from 0.0017 to 0.0034 with a mean of 0.003. Genetic similarity allowed clustering genotypes into three groups: Group 1 comprised RKO and BER, group 2 KAS-1, KAS-2, and BIS-2, group 3 consisted of HLU-1, HLU-2, and SGY. Populations displayed no signal of isolation by geographic distance (Mantel test $p>0.05$) or isolation by environmental distance (Mantel test $p>0.05$ for all environmental factors; see below). In the additional populations, 44 individual genotypes were found among the 65 sequenced, and no genotype was found at more than one site. Five accessions (RUS16, RUS17, ROD8, RUS23, RUW1) deviated from the rest and co-localized with Swedish accessions. Interestingly, we also found three individuals originating from a single site, which were virtually identical (1 SNP) to the accession TueWa1-2 from Tübingen (Germany), which was described in the Arabidopsis 1001 Genomes Project. This lineage might result from long-range migration.

Number of lines from each origin

| | |
|---|---|
| Cologne (CLN) | 44 |
| France (FRA) | 32 |
| Czechia (CZ) | 37 |
| Austria (AUT) | 11 |
| Sweden (SWE) | 39 |
| UK | 32 |
| Germany (GER) | 38 |
| Russia (RUS) | 35 |
| Netherlands (NED) | 11 |
| Spain (ESP) | 37 |
| Switzerland (SUI) | 5 |
| Italy (ITA) | 27 |
| Georgia (GEO) | 9 |

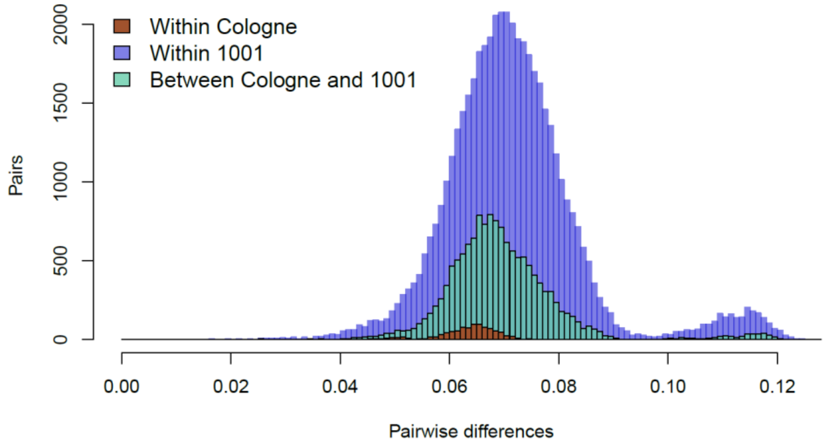


**Figure S6B:** Distribution of genetic distance between genotypes, as measured by the average number of different SNPs between each pair of genotypes divided by the total number of polymorphic positions in the dataset. Left panel shows the origin of the lines used in the study. Lines collected in Cologne (CLN) are almost as different from each other (brown) as from lines in the worldwide set (green), or as the lines of the worldwide set among each other (purple). This suggests that Cologne populations were established from a diverse pool of migrant genotypes, presumably via long range migration.

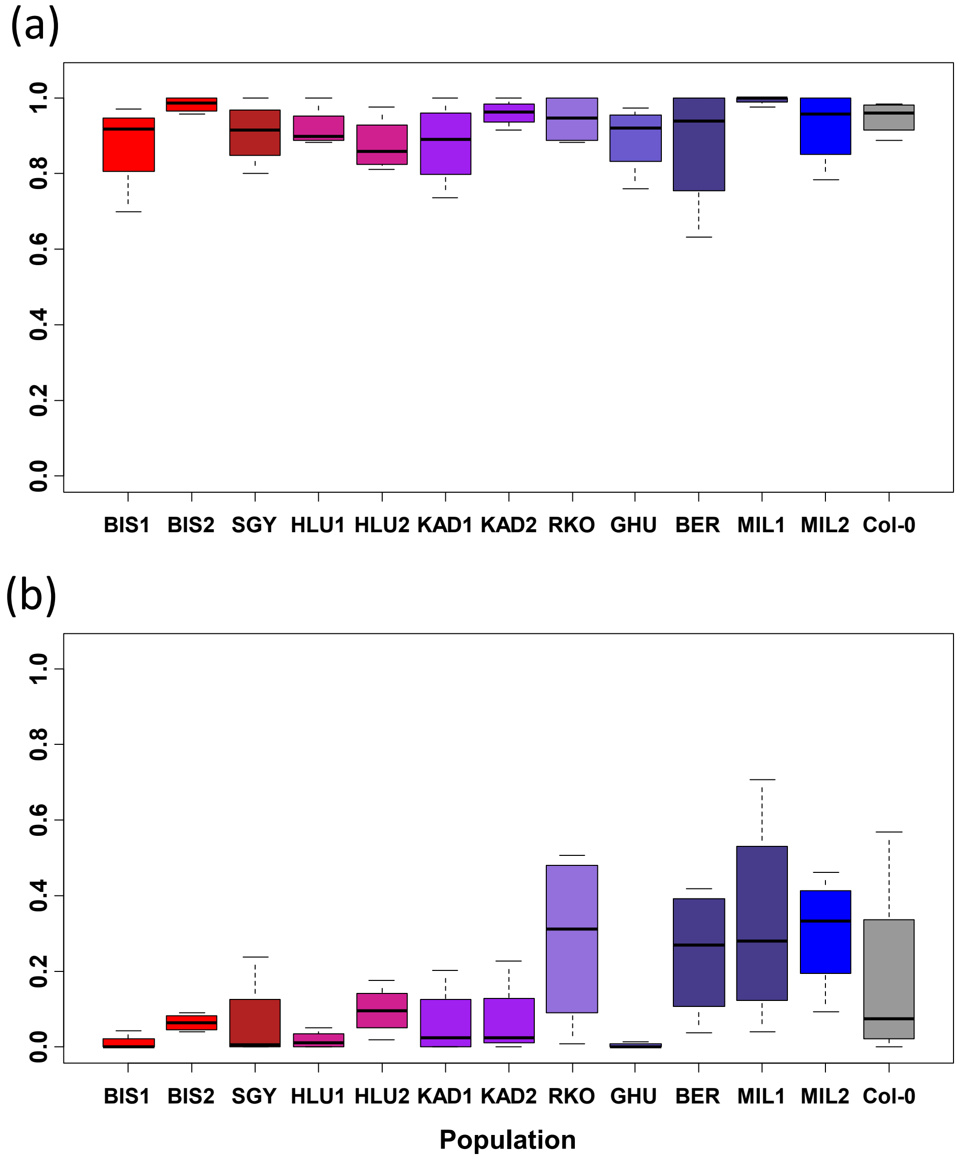


**Figure S7:** Germination frequency of seeds from the 12 different Cologne genotypes and Col-0 controls. Seeds were germinated on filter paper at 20 °C in petri dishes following stratification for 7d at 4 °C and demonstrating that seeds are viable (a) or an incubation at -21 °C in the dark (b). (4 replicates per genotype).

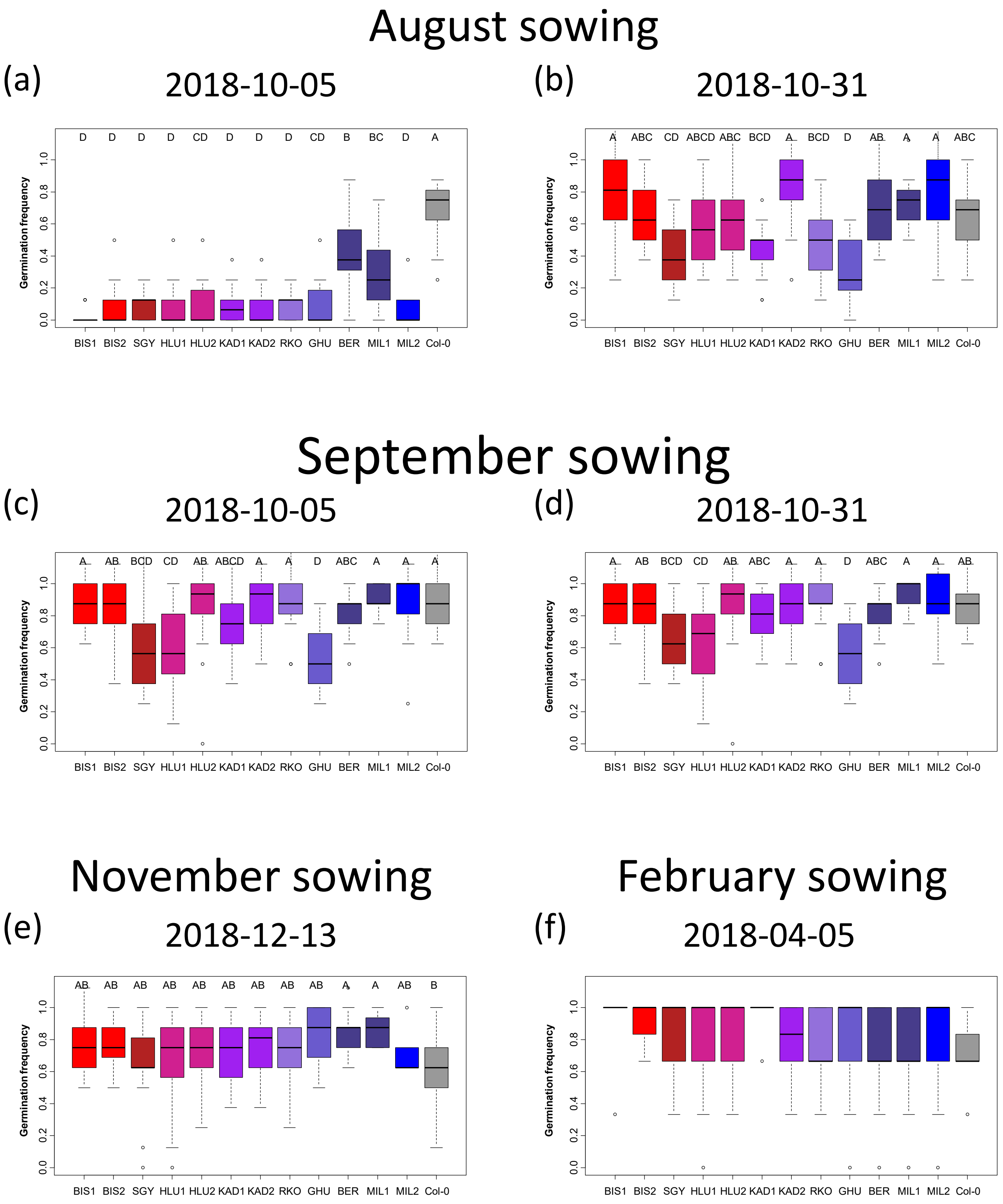


**Figure S8:** Germination frequency of seeds from the 12 different Cologne genotypes and Col-0 controls germinated in common garden conditions. Seeds were sown on dry soil on August 23rd (a, b), September 20th (c, d), November 8th (e), or February 6th (f). Germination was scored on October 5th (a, c), October 31st (b, d) for the August and September sowing. ). Note that at the second time point, more than 50% of seeds germinated, but GHU, SGY, KAS1, and RKO still had approximately 20% lower germination rates compared to the other genotypes. Germination was scored on December 13th (e), and April 5th (f) for the November and February cohorts, respectively. Differences between a) and c) can be explained by high temperature induced secondary dormancy.

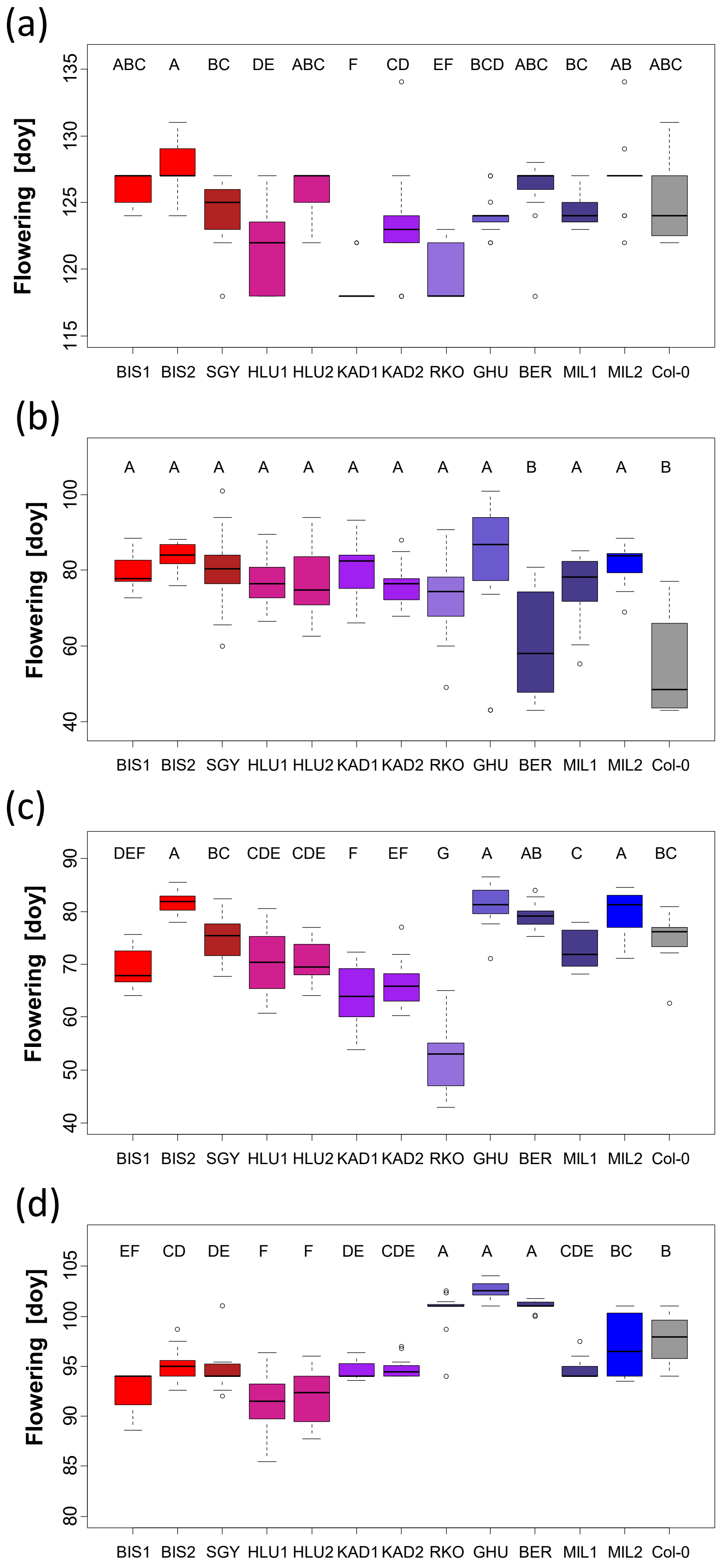


**Figure S9:** Flowering time (days of year) of plants from the 12 different Cologne genotypes and Col-0 controls grown in common garden conditions. Seeds were sown on dry soil on February 6th (a), August 23rd (b), September 20th (c), or November 8th (d). Letters indicate significant differences at a level p<0,05.

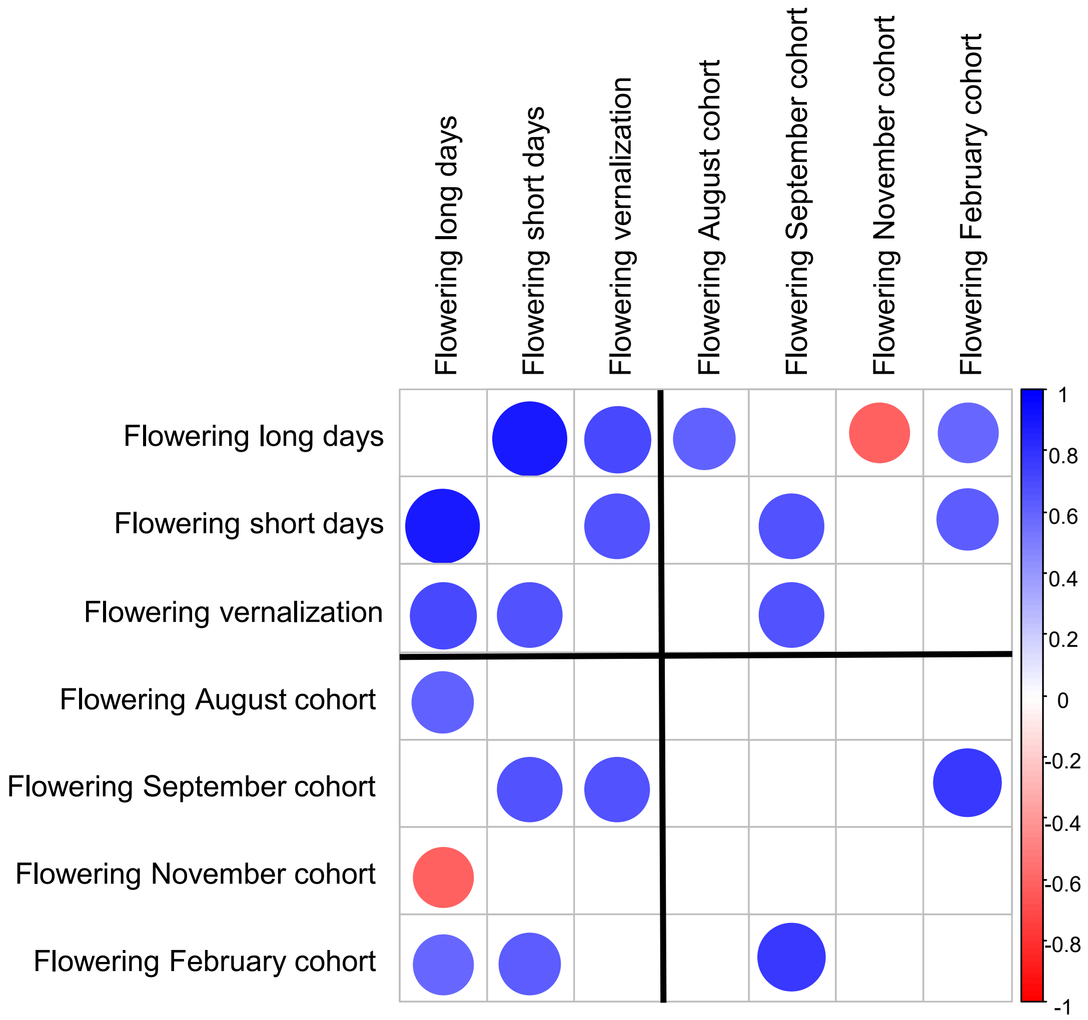


**Figure S10:** Correlation among averages for flowering time phenotypes measured in controlled laboratory conditions and in common garden experiments. Flowering times between the 4 planting cohorts show little correlation.

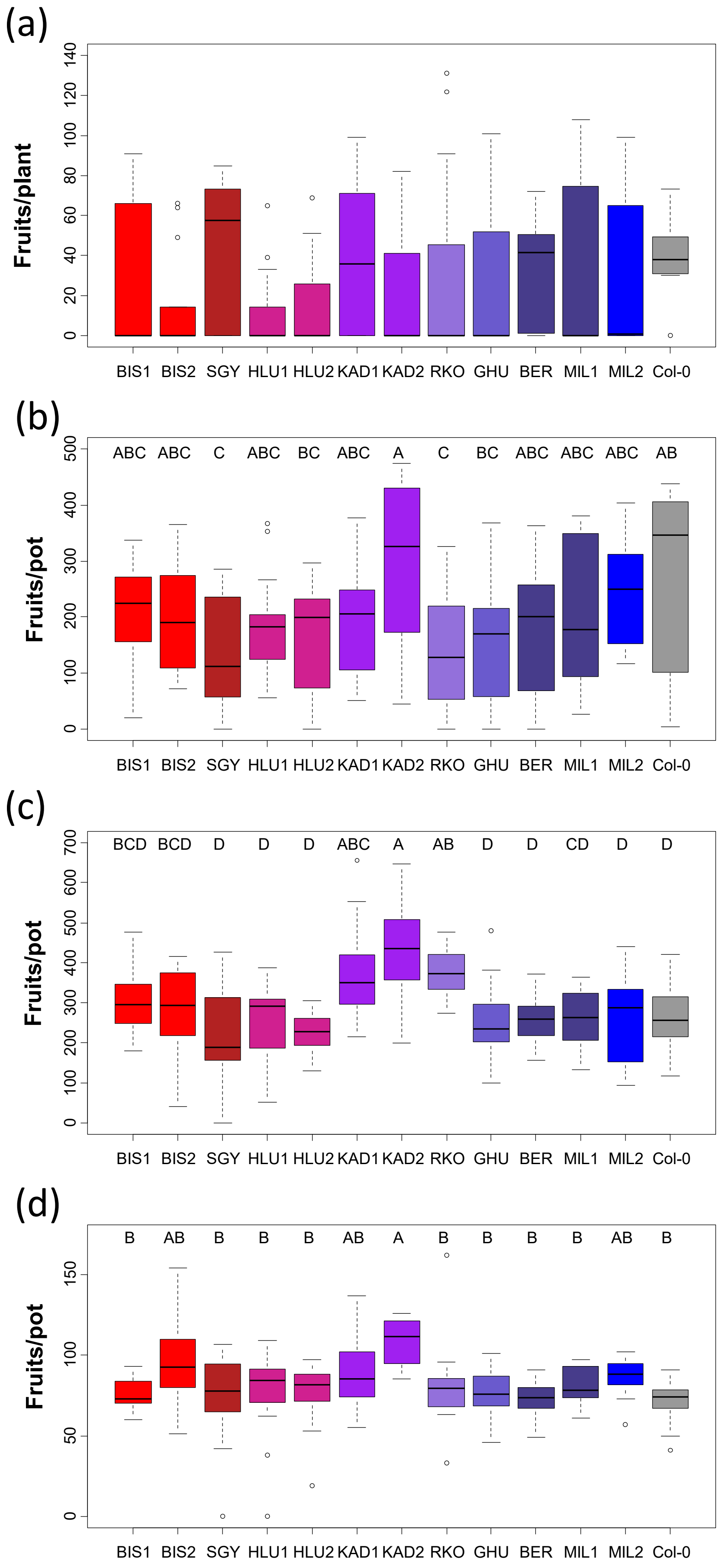


**Figure S11:** Common garden experiments - number of siliques of single plants sown in February (a), or number of siliques per pot with 8 seeds sown in August (b), September (c), or November (d). Letters indicate significant differences at a level p<0,05.Scales differ for the plots.

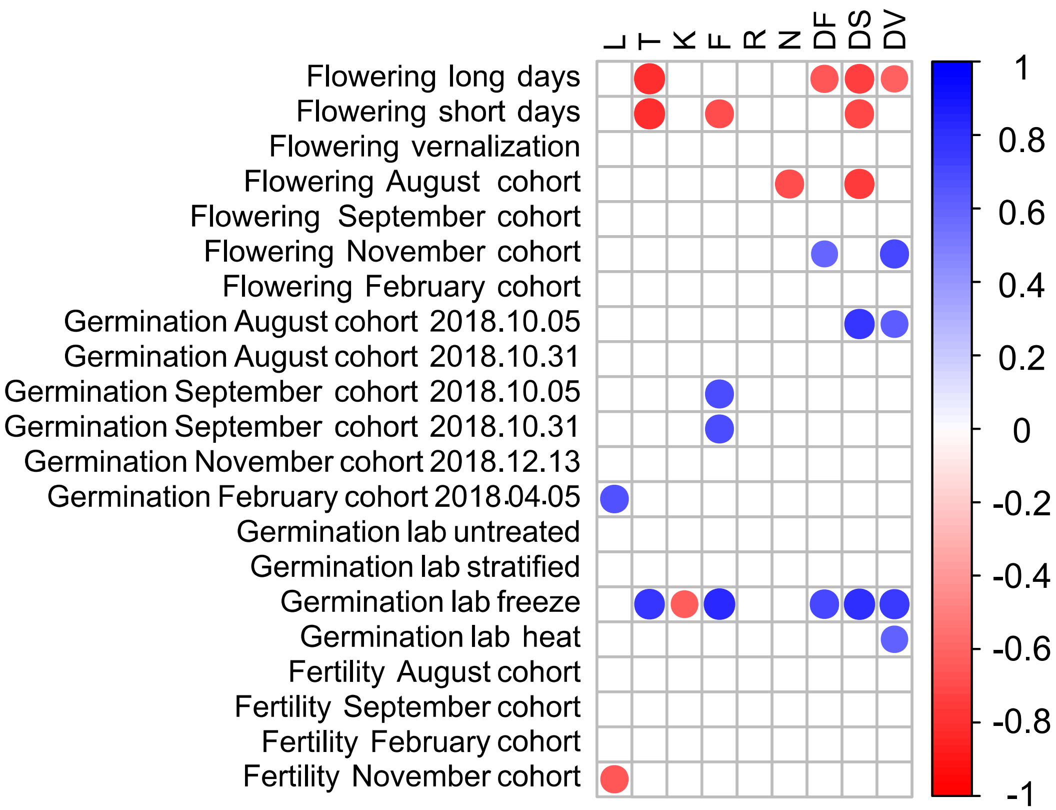


**Figure S12:** Correlation of averages of phenotypes of plants in controlled lab conditions, outdoor common garden experiments (cohorts) and averages of Ellenberg and disturbance indicator values of the sites of their origin (L- Light, T- Temperature, K- Continentality, R- pH, N- nutrients, DF- disturbance frequency, DS- disturbance severity, DV- structure-based disturbance index). Germination rates were monitored at several time points after sowing. Flowering time was scored in controlled growth chamber under long days, short days, and in a sequence combining short days, vernalization and then long days (vernalization conditions). Germination was scored in controlled conditions on untreated 2-month-old seeds and after stratification. Secondary dormancy was determined by scoring germination in stratified seeds exposed to either freezing conditions or heat (see methods for details). Fertility was measured in all four outdoor common garden cohorts, but not in growth chambers.

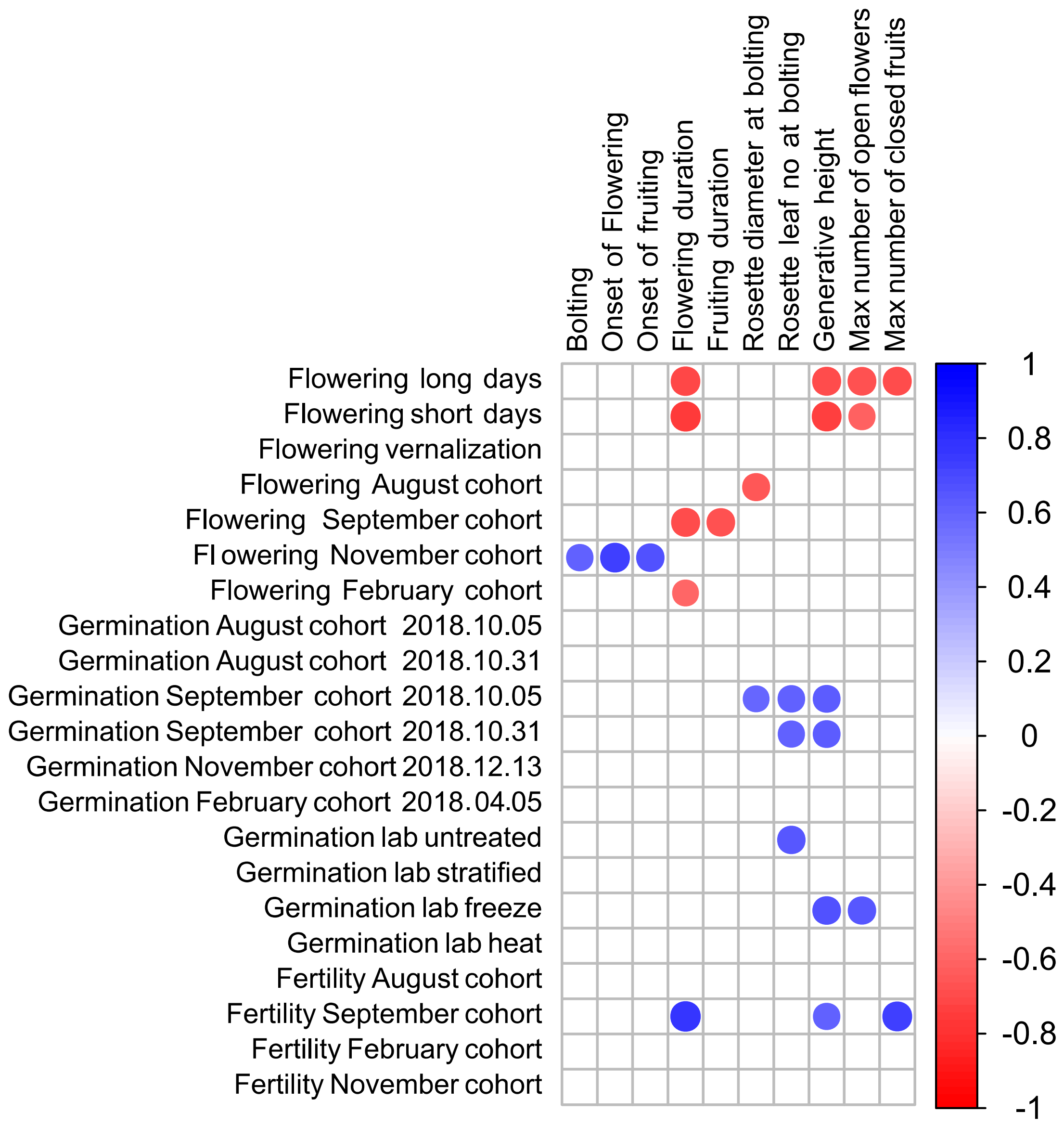


**Figure S13:** Correlation of averages of phenotypes of plants in their original sites, in controlled lab conditions and in common garden experiments. Germination rates were monitored at several time points after sowing. Flowering time was scored in controlled growth chamber under long days, short days, and in a sequence combining short days, vernalization and then long days (vernalization conditions). Germination was scored in controlled conditions on untreated 2-month-old seeds and after stratification. Secondary dormancy was determined by scoring germination in stratified seeds exposed to either freezing conditions or heat (see methods for details). Fertility was measured in all four outdoor common garden cohorts, but not in growth chambers.
